# Human erythroid progenitors express antigen presentation machinery

**DOI:** 10.1101/2024.06.27.601047

**Authors:** Rebecca L. Clements, Elizabeth A. Kennedy, David Song, Ariana Campbell, Hyun Hyung An, Kevin R. Amses, Taylor Miller-Ensminger, Mary M. Addison, Laurence C. Eisenlohr, Stella T. Chou, Kellie Ann Jurado

## Abstract

Early-life immune exposures can profoundly impact lifelong health. However, functional mechanisms underlying fetal immune development remain incomplete. Erythrocytes are not typically considered active immune mediators, primarily because erythroid precursors discard their organelles as they mature, thus losing the ability to alter gene expression in response to stimuli. Erythroid progenitors and precursors circulate in human fetuses and neonates. Although there is limited evidence that erythroid precursors are immunomodulatory, our understanding of the underlying mechanisms remains inadequate. To define the immunobiological role of fetal and perinatal erythroid progenitors and precursors, we analyzed single cell RNA-sequencing data and found that transcriptomics support erythroid progenitors as putative immune mediators. Unexpectedly, we discovered that human erythroid progenitors constitutively express Major Histocompatibility Complex (MHC) class II antigen processing and presentation machinery, which are hallmarks of specialized antigen presenting immune cells. Furthermore, we demonstrate that erythroid progenitors internalize and cleave foreign proteins into peptide antigens. Unlike conventional antigen presenting cells, erythroid progenitors express atypical costimulatory molecules and immunoregulatory cytokines that direct the development of regulatory T cells, which are critical for establishing maternal-fetal tolerance. Expression of MHC II in definitive erythroid progenitors begins during the second trimester, coinciding with the appearance of mature T cells in the fetus, and is absent in primitive progenitors. Lastly, we demonstrate physical and molecular interaction potential of erythroid progenitors and T cells in the fetal liver. Our findings shed light on a unique orchestrator of fetal immunity and provide insight into the mechanisms by which erythroid cells contribute to host defense.

## Introduction

The fetal immune system is highly specialized to meet the shifting needs of vastly different environments pre- and post-birth. *In utero*, tolerance to self-derived and maternal antigens, as well as benign food and environmental antigens, is essential to prevent harmful inflammation. The harsh postnatal environment, however, necessitates functionally mature immune cells that can respond to ever-present threats. To address these contrasting needs, fetal immunity is carefully regulated.

Failure to establish tolerance can result in fetal demise. It is therefore imperative that we gain a greater understanding of the development of immune tolerance during pregnancy. Knowledge of the human fetal and perinatal immune system has immense potential to help us understand health and disease during pregnancy, as a neonate, and likely every stage of life afterward. Deciphering molecular mechanisms of fetal immune development may hold the key to understanding evolutionarily honed paradigms of immune regulation that can be leveraged for novel immunotherapeutic approaches for transplantation, autoimmunity, and inflammatory disorders.

Regulatory T cells (T_regs_) play an essential role in mediating immune homeostasis during pregnancy^1^. Maternal-derived T_regs_ promote tolerance of paternal and fetal antigens, enabling implantation and preventing fetal rejection. Fetal-derived T_regs_ are important mediators of immunotolerance to self, maternal, and benign antigens. Fetal CD4^+^ T cells are particularly predisposed to developing into T_regs_, which is reflected in the high abundance of T_reg_ cells during gestation^1^. Aberrant T_reg_ development may lead to failure to tolerate self-derived or benign antigens, resulting in autoimmune or allergic disorders after birth. However, the molecular mechanisms that promote fetal T_reg_ development are unknown.

Major histocompatibility complex (MHC) molecules drive adaptive immune responses via presentation of antigens to CD8^+^ (MHC class I) or CD4^+^ (MHC class II) T cells. While MHC I is expressed by almost all cells, constitutive expression of MHC II is historically thought to be restricted to “professional” antigen presenting immune cells (dendritic cells, macrophages, and B cells). In contrast to this traditional dogma, several “atypical” hematopoietic and non-hematopoietic cells constitutively express MHC II; small intestinal epithelial cells^2–7^, alveolar epithelial cells^8–10^, group 3 innate lymphoid cells^11,12^, activated CD4^+^ T cells^13,14^, and hematopoietic stem cells^15^ constitutively express MHC II and associated machinery. Many other cells express MHC II in specific contexts, such as inflammatory or autoimmune disorders, or upon stimulation with interferon-gamma (IFN-γ)^2–7,16–20^. It is hypothesized that atypical antigen presenting cells mainly serve to modulate or maintain populations of antigen-experienced T cells. However, seemingly conflicting findings indicate that function and outcome of atypical antigen presentation is context-dependent. Some reports suggest that antigen presentation via atypical antigen presenting cells can lead to tolerogenic outcomes such as T_reg_ development^21–24^, anergy^13,25,26^, or apoptotic cell death^12^.

Despite close contact with other circulating immune cells, red blood cells have classically been disregarded as immunologically inactive and rather simply considered oxygen transporters. This notion emanates from the fact that erythroid cells discard their organelles during maturation in the bone marrow and thus lose their ability to modify gene expression prior to entering circulation. Nucleated erythroid progenitors and precursors circulate in developing human fetuses and neonates^27–29^. There is growing evidence that erythroid progenitors and precursors are immunomodulatory^27,28,30–45^, yet our knowledge of underlying mechanisms remains inadequate, especially in the context of human immunology.

We aimed to address this gap in knowledge regarding the immunological function of erythroid progenitors and precursors in the context of fetal and neonatal development. In this study, we describe an unexpected role for human perinatal erythroid progenitors to mediate tolerogenic adaptive immunity via MHC II-dependent antigen presentation.

## Methods

### Isolation of primary erythroid progenitors from human umbilical cord blood

Human umbilical cord blood in citrate-phosphate-dextrose solution was obtained from the Carolinas Cord Blood Bank. Umbilical cord blood was diluted 1:1 in phosphate buffered saline (PBS) and layered on top of Histopaque-1083 density medium (Sigma). Density gradient separation was performed by centrifuging at 600 x *g* for 30 minutes at room temperature without braking. Mononuclear cells were isolated and washed twice in 0.5% bovine serum albumin (BSA) in PBS at 300 x *g* for 8 minutes at room temperature. Non-specific antibody binding was blocked by incubating mononuclear cells in 10% human serum (Research Products International) and 0.5% BSA in PBS for 30 minutes on ice. Cells were stained with anti-CD235a, anti-CD71, anti-CD45, and anti-CD34 antibodies listed in Supplemental Table 1 for 30 minutes on ice, stained with 1 µM Hoechst 33342 (AAT Bioquest) for 5 minutes on ice, washed twice in 0.5% BSA in PBS at 300 *x g* for 5 minutes, and resuspended in 0.5% BSA in PBS. For flow cytometry-based analyses, erythroid cells were identified via gating (not isolation). For all other experiments, erythroid cells were isolated via fluorescence activated cell sorting. Cells were sorted on a FACSAria II Cell Sorter (BD Biosciences). Erythroid progenitors are defined as CD235a^+^CD71^+^Hoechst 33342^+^CD45^+^. Erythroid precursors are defined as CD235a^+^CD71^+^Hoechst 33342^+^CD45^-^. Reticulocytes are defined as CD235a^+^CD71^+^Hoechst 33342^-^CD45^-^. Isolated cells were assayed immediately or cultured overnight in StemSpan SFEM (STEMCELL) with 50 ng/mL recombinant human stem cell factor (PeproTech), 10^-6^ M dexamethasone (Sigma), and 2 U/mL recombinant human erythropoietin (PeproTech) in a humidified, 37°C incubator at 5% CO_2_.

### Direct-quenched ovalbumin antigen processing assay

100,000 unstimulated, undifferentiated HUDEP-2 or THP-1 cells were incubated with 10 μg/mL DQ-ovalbumin (Invitrogen) at 4°C for 4 hr or 37°C for 4 hr, 2 hr, or 1 hr. Cells were washed three times in 0.5% BSA in PBS and fluorescence was measured via flow cytometry. Primary umbilical cord blood mononuclear cells were isolated via density gradient as described above. 1,000,000 primary cord blood mononuclear cells were incubated with 10 μg/mL DQ-ovalbumin at 37°C for 3 hr. Cells were subsequently stained for flow cytometry with anti-CD235a, anti-CD71, anti-CD34, and Hoechst 33342. Fluorescence of DQ-ovalbumin was measured via flow cytometry. Distinct populations of erythroid cells were identified and gated as described above. Induced pluripotent stem cell-derived red blood cells (iRBCs) were differentiated as we performed previously^46^ (see Supplemental Methods). After 3 days of differentiation, iRBCs were isolated via magnetic activated cell sorting of anti-CD235a magnetic beads. 100,000 iRBCs were incubated with 10 μg/mL DQ-ovalbumin at 4°C or 37°C for 4 hr. Cells were washed three times in 0.5% BSA in PBS and fluorescence was measured via flow cytometry.

### Analysis of single cell RNA-sequencing datasets

Single cell RNA-sequencing data was obtained from Gene Expression Omnibus (GEO) accession number GSE150774 (umbilical cord blood)^47^, GSE160251 (second trimester fetal liver)^48^, or ArrayExpress accession E-MTAB-11343 (fetal development)^49^. Data were analyzed using R package Seurat^50^.

### Bulk RNA-sequencing analysis

Three days post-differentiation, iRBCs were isolated via magnetic sorting. 1×10^6^ differentiated iRBCs or undifferentiated HUDEP-2 cells were stimulated with 50 ng/mL recombinant human interferon-gamma (PeproTech) for 6 hours. Stimulated cells and unstimulated controls were resuspended in TRIzol (Invitrogen) and RNA was isolated via phase separation using 1-bromo-3-chloropropane. Aqueous layer was precipitated in isopropanol and RNA was washed in 70% ice-cold ethanol, then resuspended in nuclease-free water. Library prep was performed using stranded mRNA prep with poly-A enrichment (Illumina). Sequencing was performed at the Children’s Hospital of Philadelphia High-Throughput Sequencing Core using NextSeq 1000 (Illumina). Raw sequencing read counts were normalized and processed using R.

### Cell-cell interaction analysis

CellPhoneDB^51^ was used to infer cell-cell interactions from single cell RNA-sequencing data^49^ using default parameters.

## Results

### Human erythroid progenitors express MHC II antigen presentation machinery

To identify immunological features of erythroid progenitors, we queried publicly available transcriptomic data. We analyzed single-cell RNA-sequencing data of human umbilical cord blood erythroid cells designed to find regulators of erythropoiesis^47^. We filtered this dataset to remove non-erythroid contaminants, then re-clustered the data (Figure 1A) and identified clusters via surface markers and transcription factors differentially expressed throughout erythropoiesis (Supplemental Figure 1; Figure 1C). Committed erythroid progenitors (BFU-E, CFU-E, and proerythroblasts) are transcriptionally distinct from erythroid precursors^52^. Surprisingly, we found that erythroid progenitors express antigen presentation machinery that is typically restricted to professional antigen presenting immune cells. In our analysis, we found that erythroid progenitors constitutively express HLA-DR, DP, and DQ alpha and beta chains at moderate levels (Figure 1D). Expression of MHC II is diminished in basophilic erythroblasts and subsequent stages of erythropoiesis. Consistent with our transcriptomic analysis, flow cytometry of primary erythroid progenitors from human umbilical cord blood from eight independent donors confirmed constitutive surface expression of HLA-DR protein in erythroid progenitors (CD45^+^), but not erythroid precursors (CD45^-^) or enucleated reticulocytes (Figure 1F-G). Previous studies reported that murine erythroid progenitors do not express MHC II RNA^15^. To confirm this, we assessed MHC II (I-A/I-E) protein expression via flow cytometry in erythroid progenitors isolated from neonatal spleen and adult bone marrow. We do not observe I-A/I-E surface expression in adult or neonatal murine erythroid progenitors, even upon stimulation with IFN-γ (Supplemental Figure 2). These data indicate species-specific expression of MHC II antigen presentation machinery in erythroid progenitors.

**Figure 1.**
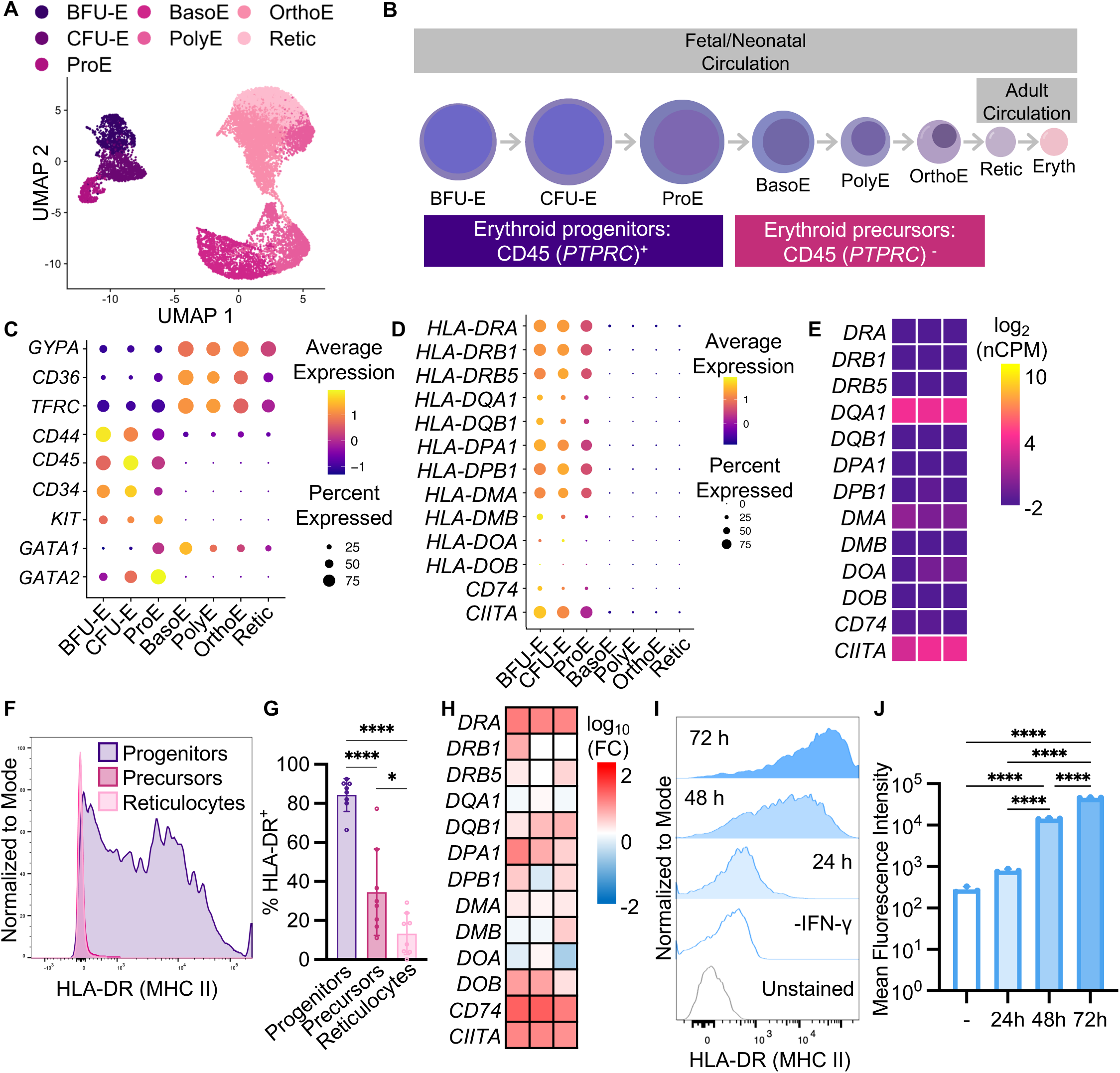
Human erythroid progenitors express MHC class II antigen presentation machinery. **A)** UMAP plot of single-cell RNA-sequencing data of primary human umbilical cord erythroid cells. Data from Huang, *et al.* (GSE150774). **B)** Schematic of erythropoiesis. **C)** Analysis parameters and clustering of single-cell RNA-sequencing data was validated with known surface markers of erythropoiesis and transcription factors differentially expressed during erythropoiesis. For clarity, common protein names shown instead of gene names. *GYPA* = CD235a, *TRFC* = CD71, *PTPRC* = CD45, *KIT* = CD117. **D)** Single cell RNA-sequencing of primary cord blood erythroid progenitors reveals expression of MHC class II and associated machinery in erythroid progenitors. **E)** Bulk RNA-sequencing of unstimulated, undifferentiated HUDEP-2 cells demonstrates low constitutive expression of MHC class II and associated machinery. Heatmap displays normalized counts per million (nCPM) for three replicates. **F)** Flow cytometry of primary cord blood erythroid progenitors indicates surface expression of MHC class II (HLA-DR) in erythroid progenitors (CD45^+^) but not precursors (CD45^-^) or reticulocytes. **G)** Quantification of (F). Graph indicates percentage of HLA-DR- positive erythroid cells of eight independent donors. Mean ± SD. One-way ANOVA. *P<0.0332, ****P < 0.0001. **H)** Bulk RNA-sequencing of undifferentiated HUDEP-2 cells reveals that erythroid progenitors upregulate expression of MHC class II molecules and associated machinery in response to stimulation with interferon-gamma (IFN-γ, 50 ng/mL, 6h). Heatmap displays fold-change (FC) compared to unstimulated control for three replicates. **I)** Flow cytometry of undifferentiated HUDEP-2 cells reveals that erythroid progenitors upregulate surface expression of MHC Class II (HLA- DR) in response to stimulation with interferon-gamma (IFN-γ, 50 ng/mL). **J)** Quantification of (I). Graph indicates mean fluorescence intensity. Mean ± SD. One-way ANOVA. ****P < 0.0001. n = 3 technical replicates per experiment, representative of 3 independent experiments.

To better understand kinetics of MHC II expression, we utilized immortalized <u>H</u>uman <u>U</u>mbilical cord blood-<u>D</u>erived <u>E</u>rythroid <u>P</u>rogenitor (HUDEP-2) cells^53^. HUDEP-2 cells are more tractable and uniformly differentiated compared to umbilical cord blood-derived erythroid progenitors. Using bulk RNA-sequencing, we find that HUDEP-2 cells express MHC II RNA, albeit at low levels (Figure 1E). Undifferentiated HUDEP-2 cells are primarily BFU-E, CFU-E, and proerythroblasts (Supplemental Figure 3A). Six days post-differentiation, HUDEP-2 cells are composed mostly of basophilic erythroblasts and polychromatic erythroblasts (Supplemental Figure 3B). Using flow cytometry, we find that undifferentiated HUDEP-2 cells constitutively express low levels of HLA-DR protein on the cell surface (Figure 1I-J). However, six days post-differentiation, HUDEP-2 cells no longer express HLA-DR protein, consistent with our transcriptomic data (Supplemental Figure 4). These data indicate that MHC II expression is unique to erythroid progenitors and is diminished in later stages of erythropoiesis.

Antigen presentation via MHC II requires successful peptide loading and trafficking of the MHC-peptide complex. The invariant chain (CD74) and its cleavage fragment, class II-associated invariant chain peptide (CLIP), temporarily occupy the peptide-binding groove of MHC II to prevent peptide binding in the ER^54–56^. The molecular chaperone HLA-DM facilitates the exchange of CLIP for antigenic peptide in the endo-lysosome^54^. In our transcriptomic analysis, we found that erythroid progenitors constitutively express CD74 and HLA-DM (Figure 1D). We found that HLA-DO, which regulates HLA-DM activity^57–59^, has very low expression levels in erythroid progenitors (Figure 1D). Thus, erythroid progenitors express machinery required for peptide loading onto MHC II.

### Erythroid progenitors upregulate MHC II expression in response to interferon-gamma

Class II transactivator (CIITA) is a non-DNA binding co-activator that serves as the sole master regulator of MHC II gene expression^60^. Production of CIITA is both tightly controlled and highly specific for MHC II-producing cells. Expression of MHC II and associated machinery including the invariant chain and HLA-DM is controlled by CIITA^60^. In our transcriptomic analysis, we found that erythroid progenitors constitutively express CIITA (Figure 1D-E). In humans, four different promoters (pI-pIV) drive CIITA transcription^61–63^. Using qPCR primers specific for the unique first exon of each promoter, we find that undifferentiated HUDEP-2 cells utilize primarily pI at baseline, the so-called myeloid-specific promoter (Supplemental Figure 5)^61,63^. Interferon-γ induces CIITA transcription, and ultimately upregulates MHC II expression^60^. In our transcriptomic analysis, we found that erythroid progenitors express the IFN-γ receptor and are thus capable of responding to IFN-γ stimulation.

Using undifferentiated HUDEP-2 cells, we demonstrate that erythroid progenitors upregulate both RNA and protein expression of MHC II genes in response to IFN-γ. Upon IFN-γ stimulation, undifferentiated HUDEP-2 cells upregulate expression of CIITA via pIV and pIII promoters (Supplemental Figure 5). In addition to upregulating transcription of CIITA, HUDEP-2 cells also upregulate transcription of HLA-DR, DP, and DQ alpha and beta chains, the invariant chain, and HLA-DM upon IFN-γ stimulation (Figure 1H). Consistent with RNA upregulation, we found that undifferentiated HUDEP-2 cells increase surface expression of HLA-DR protein upon IFN-γ stimulation in a time-dependent manner (Figure 1I-J). While unstimulated HUDEP-2 cells express relatively low levels of HLA-DR on the cell surface, after 72 hours of IFN-γ stimulation, HUDEP-2 cells express HLA-DR at levels higher than activated macrophage-like THP-1 cells (Supplemental Figure 6)^64^. These findings suggest that erythroid progenitors not only express MHC II machinery at baseline, but also increase expression upon activation with inflammatory signals.

### Erythroid progenitors process exogenous antigens via proteolytic cleavage

Classical MHC II antigen processing relies on internalization of exogenous proteins followed by proteolytic cleavage^54^. To determine if erythroid progenitors are capable of internalizing exogenous proteins, we incubated HUDEP-2 cells with ovalbumin conjugated to Alexa Fluor 488 fluorescent dye (OVA-AF488). Undifferentiated HUDEP-2 cells were capable of internalizing OVA-AF488, as measured via flow cytometry (Supplemental Figure 7). However, differentiated HUDEP-2 cells were unable to internalize OVA-488 (Supplemental Figure 7). This indicates that erythroid progenitors possess endocytic capabilities that are lost in erythroid precursors. After internalization, foreign proteins are cleaved in the endo-lysosome. In our single cell transcriptomic analysis, we found that erythroid progenitors express a variety of endo-lysosomal proteases important for antigen processing including several cathepsin proteases (Figure 2A)^65^. We demonstrate functional cathepsin S and cathepsin D protease activity in HUDEP-2 cell lysates, and find that activity levels are similar to those found in the monocytic THP-1 cell line (Supplemental Figure 8)^64^.

**Figure 2.**
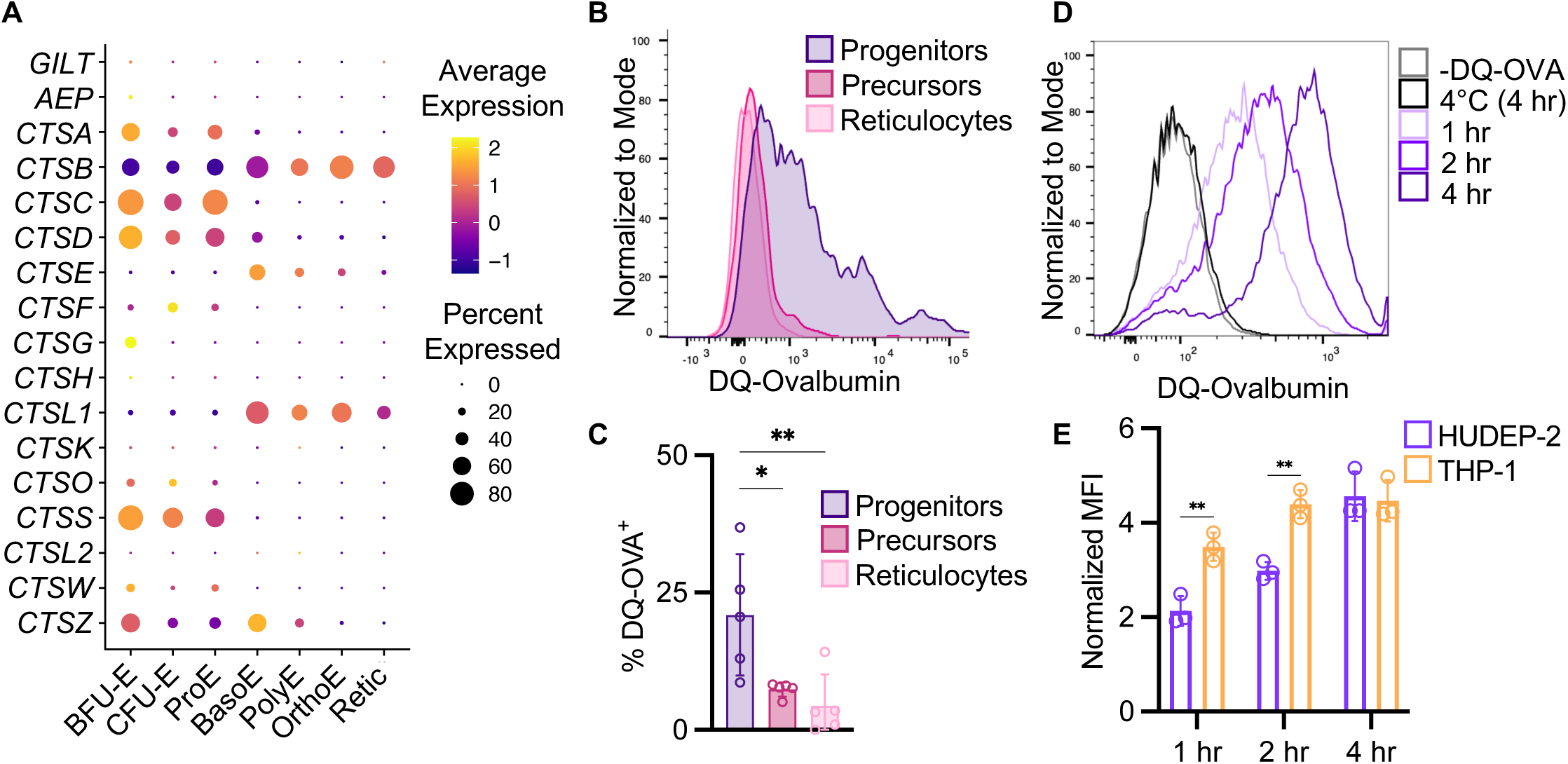
Human erythroid progenitors process antigens. **A)** Human erythroid progenitors express antigen processing machinery. Single cell RNA-sequencing of primary cord blood erythroid cells reveals expression of endo- lysosomal proteases important for antigen uptake and cleavage. Data from Huang, *et al.* (GSE150774). **B)** Erythroid progenitors internalize and cleave antigens. Primary human umbilical cord blood mononuclear cells were incubated with 10 μg/mL direct quenched-ovalbumin (DQ-OVA) at 37°C for 3 hr. Fluorescence measured via flow cytometry indicates relief of self-quenching & successful receptor-mediated endocytosis/proteolytic digestion. Erythroid progenitors were queried for their internalization and processing of DQ-OVA after gating on either erythroid progenitors (CD34+), precursors (CD34-), or reticulocytes. **C)** Quantification of (B). Graph indicates percentage of HLA-DR-positive erythroid cells of five independent donors. Mean ± SD. One-way ANOVA. *P<0.03, **P < 0.002. **D)** Undifferentiated, unstimulated HUDEP-2 cells or THP-1 cells were incubated with 10 μg/mL DQ-OVA at 4°C for 4 hr or 37°C for indicated times. **E)** Quantification of (D). Graph indicates mean fluorescence intensity normalized to -DQ-OVA control. Mean ± SD. Two-way ANOVA (only comparisons between HUDEP-2 and THP-1 shown on graph). **P < 0.002. n = 3 technical replicates per experiment, representative of 3 independent experiments.

Given our data demonstrating that erythroid progenitors internalize antigen and express endo-lysosomal proteases, we next aimed to determine whether erythroid progenitors are capable of processing antigens via proteolytic cleavage. To test this, we utilized direct quenched ovalbumin (DQ-OVA), ovalbumin protein conjugated to a self-quenching dye. Upon internalization and proteolytic cleavage of DQ-OVA, self-quenching is released and the fragments of DQ-OVA fluoresce. In primary erythroid progenitors from human umbilical cord blood, we find that erythroid progenitors, but not erythroid precursors or reticulocytes, are capable of processing DQ-OVA (Figure 2B-C). These data confirm that erythroid progenitors can internalize and cleave synthetic antigens. Using undifferentiated HUDEP-2 cells, we find that this ability is temperature-dependent, as HUDEP-2 cells are unable to process DQ-OVA at 4°C (Figure 2D). While we observe a time-dependent increase in DQ-OVA processing, we find that in comparison to the monocytic cell line THP-1, undifferentiated HUDEP-2 cells are less efficient at processing antigen. We see a significant difference between THP-1 and HUDEP-2 mean fluorescence at 1 and 2 hours, and this difference is abrogated by 4 hours (Figure 2E; Supplemental Figure 9). Consistent with our observations that erythroid precursors are unable to internalize OVA-488, we find that differentiated HUDEP-2 cells do not process DQ-OVA (Supplemental Figure 9A-B). Together, these data indicate that erythroid progenitors are capable of processing exogenous antigens via proteolytic cleavage.

### Erythroid progenitors lack classical costimulatory molecules

T cell survival, differentiation, and effector functions are primarily regulated by costimulatory molecules on antigen presenting cells binding to their associated receptors on T cells^66^. T cell activation via MHC II requires costimulation in addition to antigen-specific recognition via the T cell receptor^67^. In our transcriptomic analysis, we see negligible expression of classical costimulatory molecules *CD80*, *CD86*, and *ICOSL* in umbilical cord blood erythroid progenitors and HUDEP-2 cells (Figure 3A-B). We confirmed that primary umbilical cord erythroid progenitors lack classical costimulatory molecules on their cell surface via flow cytometry. Furthermore, expression of classical costimulatory molecule RNA or protein is not induced upon stimulation with IFN-γ in undifferentiated HUDEP-2 cells (Figure 3E; Supplemental Figure 11). Several non-classical costimulatory molecules can interact with T cell co-receptors to influence T cell fate^66^. Our single cell transcriptomic analysis indicated expression of several non-classical costimulatory molecules in erythroid progenitors including HVEM, CD48, Gal9, and VISTA (Figure 3A). HVEM is expressed by several hematopoietic and non-hematopoietic cells and can act as either a costimulatory molecule or co-inhibitory molecule depending on the co-receptors present on the T cell surface^68^. We confirmed surface expression of HVEM on erythroid progenitors via flow cytometry (Figure 3C-D). Stimulation of undifferentiated HUDEP-2 cells with IFN-γ upregulates surface expression of HVEM and influences RNA expression patterns in erythroid progenitors (Figure 3E-G). While our single cell transcriptomics analysis demonstrated expression of VISTA and Gal9 RNA, we do not observe VISTA or Gal9 surface expression via flow cytometry, even upon stimulation with IFN-γ (Supplemental Figure 11). This is consistent with previous findings that cord blood erythroid cells express VISTA RNA but do not express VISTA protein on the cell surface^33^. Overall, consistent with other atypical antigen presenting cells, we find that erythroid progenitors lack classical costimulatory molecules and instead express non-classical and co-inhibitory molecules.

**Figure 3.**
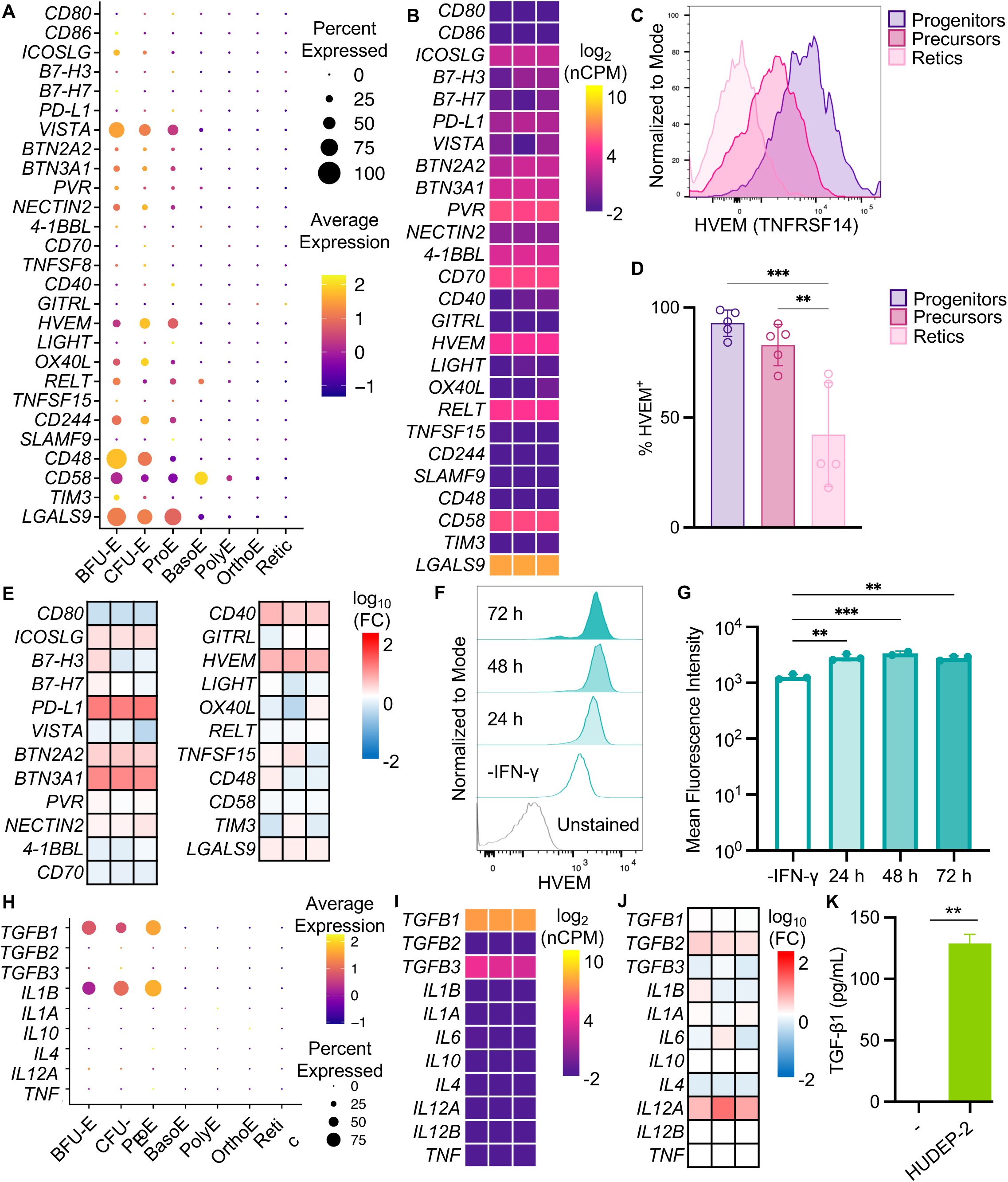
Human erythroid progenitors lack classical co-stimulation machinery and secrete immunosuppressive cytokines. **A)** Single cell RNA-sequencing of primary human cord blood erythroid progenitors reveals expression of non-classical co-stimulatory molecules in erythroid progenitors. For clarity, common protein names shown instead of gene names. *CD276* = B7-H3, *HHLA2* = B7-H7, *CD274* = PD-L1, *C10orf54* = VISTA, *PVR* = CD155, *NECTIN2* = CD112, *TNFSF9* = 4-1BBL, *TNFSF8* = CD153, *TNFSF18* = GITRL, *TNFRSF14* = HVEM, *TNFSF14* = LIGHT, *TNFSF4* = OX40L, *TNFSF15* = TL1A, *CD244* = SLAMF4, *HAVCR2* = TIM-3, *LGALS9* = Gal9. **B)** Bulk RNA-sequencing of unstimulated, undifferentiated HUDEP-2 cells demonstrates lack of classical co-stimulatory molecules. Heatmap displays normalized counts per million (nCPM) for three replicates. **C)** Flow cytometry of primary cord blood erythroid progenitors indicates surface expression of HVEM in erythroid progenitors (CD45^+^) but not precursors (CD45^-^) or reticulocytes. **D)** Quantification of (C). Graph indicates percentage of HVEM-positive erythroid cells of five independent donors. Mean ± SD. One-way ANOVA. **P < 0.02, ***P < 0.0002. **E)** Bulk RNA-sequencing of undifferentiated HUDEP-2 cells reveals that erythroid progenitors alter expression of co-stimulatory molecules in response to stimulation with interferon-gamma (IFN-γ, 50 ng/mL, 6h). Heatmap displays fold-change (FC) compared to unstimulated control for three replicates. **F)** Flow cytometry of undifferentiated HUDEP-2 cells reveals that erythroid progenitors upregulate surface expression of HVEM in response to stimulation with interferon-gamma (IFN-γ, 50 ng/mL). **G)** Quantification of (F). Graph indicates mean fluorescence intensity. Mean ± SD. One-way ANOVA. **P < 0.002, ***P < 0.0002. **H)** Single cell RNA-sequencing of primary human cord blood erythroid progenitors reveals expression of instructive cytokines in erythroid progenitors. **I)** Bulk RNA-sequencing of unstimulated, undifferentiated HUDEP-2 cells demonstrates expression of instructive cytokines. Heatmap displays normalized counts per million (nCPM) for three replicates **J)** Bulk RNA-sequencing of undifferentiated HUDEP-2 cells reveals that erythroid progenitors alter expression of instructive cytokines in response to stimulation with interferon-gamma (IFN-γ, 50 ng/mL, 6h). Heatmap displays fold-change (FC) compared to unstimulated control for three replicates. **K)** ELISA of HUDEP-2 conditioned media or non-conditioned media control indicates secretion of TGF-β1 by erythroid progenitors. Mean ± SD. Unpaired, two-tailed t-test. **P < 0.002 n = 3 technical replicates per experiment, representative of 3 independent experiments.

### Erythroid progenitors express immunoregulatory cytokines

Antigen presenting cells further influence T cell differentiation via secretion of instructive cytokines^69^. Transforming growth factor beta 1 (TGF-β1) is an immunosuppressive cytokine that inhibits proliferation of activated T cells. *TGFB1* RNA expression has been described previously in human cord blood and placental erythroid cells^32^. In our single cell transcriptomics analysis of primary cord blood erythroid cells, in line with previous findings, we observe expression of *TGFB1* RNA (Figure 3H). In addition, we also observe expression of interleukin 1 beta (IL-1β, Figure 3H), a proinflammatory instructive cytokine^69^. Our bulk transcriptomic analysis of undifferentiated HUDEP-2 cells confirmed that *TGFB1* RNA is expressed in unstimulated erythroid progenitors, but we find that *IL1B* is not produced, even upon stimulation with IFN-γ (Figure 3I-J). We confirmed production and secretion of TGF-β1 protein via ELISA of HUDEP-2 culture supernatants (Figure 3K). Overall, these findings suggest that erythroid progenitors produce instructive cytokines that could putatively influence T cell differentiation and proliferation.

### Expression of antigen processing and presentation machinery increases throughout fetal development

Erythropoiesis happens in several waves during development. “Primitive” erythropoiesis is initiated in the yolk sac, giving rise to the earliest erythroid precursors^70^. The next wave of erythroid cells arises from erythro-myeloid progenitors, which are formed in the yolk sac before seeding the fetal liver^70^. The final wave arises from hematopoietic stem cells (HSCs) derived from hemogenic endothelium in the aorta-gonad-mesonephros region beginning at 4-6 weeks post-conception^70^. These HSCs first seed the liver before engrafting in the bone marrow. HUDEP-2 cells and primary erythroid progenitors isolated from umbilical cord blood are both representative of perinatal erythroid progenitors. To better understand the role of erythroid progenitors throughout fetal development, we utilized induced pluripotent stem cell (iPSC)-derived erythroid progenitors that resemble embryonic erythroid progenitors from 4-6 post-conception week (PCW) human fetuses (iRBCs, Supplemental Figure 12)^71^. We observe that iRBCs constitutively express low levels of MHC II RNA and protein and are less responsive to IFN-γ stimulation than HUDEP-2 cells (Supplemental Figure 13). Furthermore, iRBCs are unable to process DQ-OVA (Supplemental Figure 10C-D).

We hypothesized that the differences in MHC II expression we observe between iRBCs and perinatal erythroid progenitors are due to differences in the developmental state of each model (embryonic versus perinatal). To understand whether erythroid progenitors alter expression of antigen presentation machinery throughout fetal development, we analyzed a publicly available single cell transcriptomics dataset from 7-17 PCWs^49^. T cell progenitors populate the fetal thymus after ∼9 PCWs and mature T cells are first observed ∼12 PCWs^72^. We therefore pooled data from 7-9 PCW fetuses (mid-first trimester), 10-12 PCW fetuses (late first trimester), or 13-17 PCW (early second trimester), corresponding to these milestones in T cell development (Supplemental Figure 14). We observed an age-dependent increase in expression of MHC II machinery, cathepsin proteases, costimulatory molecules, and instructive cytokines in erythroid progenitors corresponding to the appearance of T cell progenitors and mature T cells (Supplemental Figure 15). We see a lack of MHC II machinery in mid-first trimester, low expression in late first trimester, and moderate expression in early second trimester erythroid progenitors (Figure 4; Supplemental Figure 15). In a separate mid-second trimester (21 PCW) fetal liver transcriptomics dataset^70^, we see moderate expression of MHC II machinery in erythroid progenitors, reflecting our observations from full term umbilical cord blood (Supplemental Figure 16).

**Figure 4.**
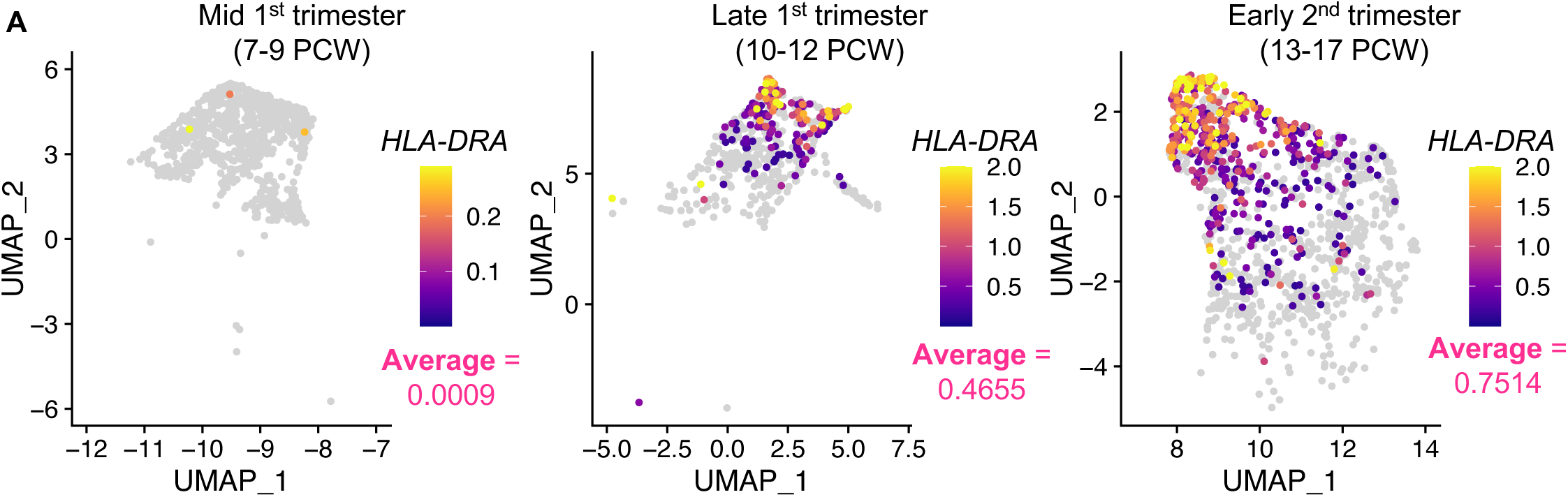
Erythroid progenitors do not express MHC class II machinery until 10 weeks post-conception. **A)** Single cell RNA-sequencing of human fetal erythroid progenitors indicates upregulation of MHC class II machinery throughout fetal development (Data from Suo, *et al.*). Erythroid progenitor cluster (BFU-E, CFU-E, and ProE) shown with *HLA-DRA* expression for each cell and the average expression levels. PCW = post-conception weeks. Data from Suo, *et al*.

### Erythroid progenitors putatively interact with fetal T cells in utero

Given that erythroid progenitors express atypical costimulatory molecules, we inquired whether fetal T cells express compatible atypical receptors. We utilized CellPhoneDB^51^ to interrogate cell-cell interactions using single cell RNA-sequencing data from second trimester fetuses^49^. Our findings demonstrate that MHC II^+^ erythroid progenitors can interact bidirectionally with T cells, with the highest number of significant (p<0.05) ligand-receptor pairs occurring with T_reg_ cells, followed by unconventional type 3 innate T cells (Figure 5; Supplemental Figure 17). We identified ligand-receptor pairs that function in costimulation, chemokine signaling, cell-cell adhesion, and regulation of T cell differentiation, proliferation, and survival. This confirmation that erythroid progenitors and fetal T cells express complementary ligand-receptor pairs underscores their potential to interact.

**Figure 5.**
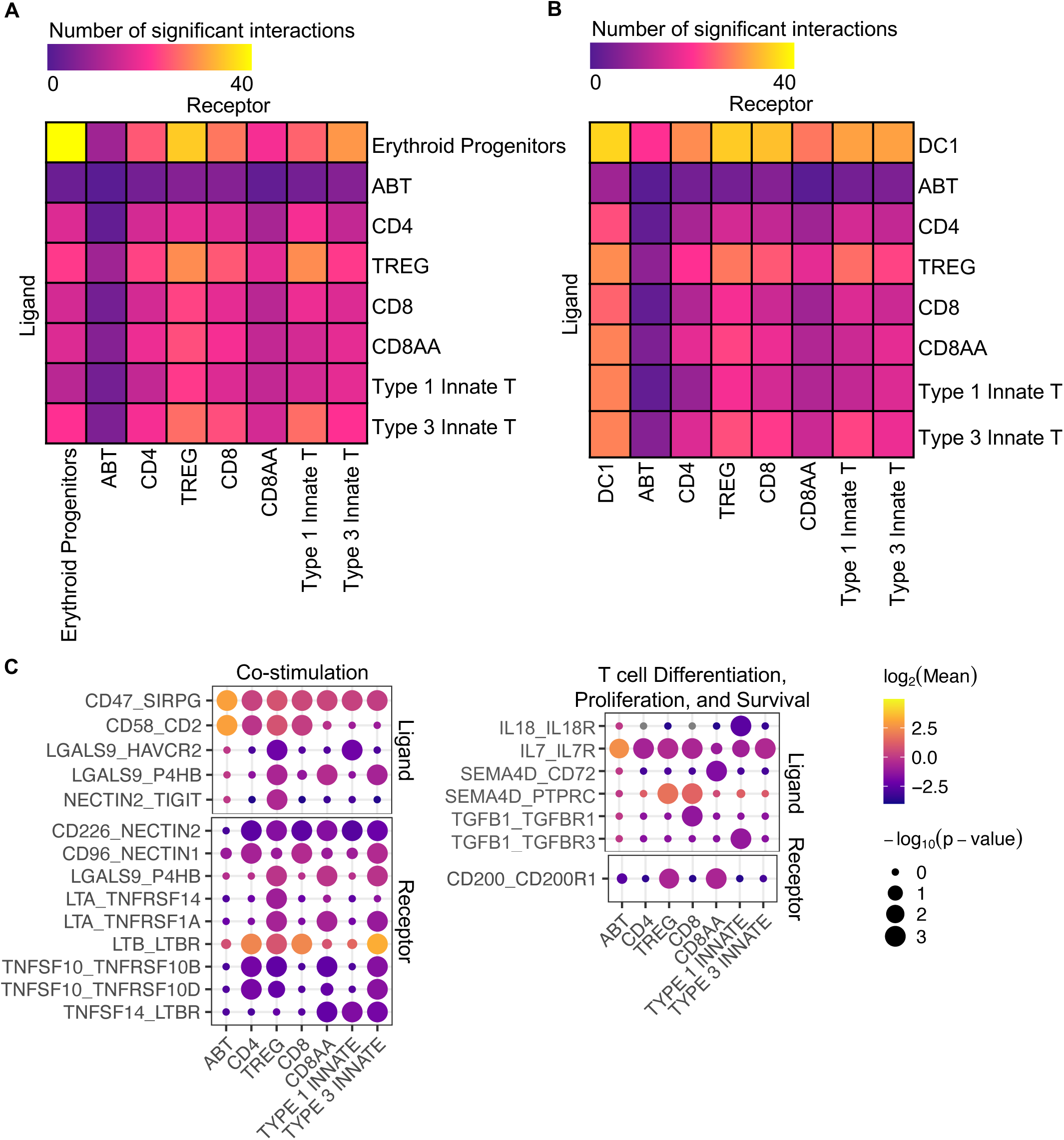
Erythroid progenitors and fetal T cells express complementary ligand-receptor pairs. Cell-cell interactions were inferred using single cell RNA-sequencing of T cells and **A)** MHC II^+^ erythroid progenitors or **B)** type 1 dendritic cells (Data from Suo, *et al.*). Significant ligand-receptor pairs (p<0.05) were analyzed using CellPhoneDB. For each significant interaction, erythroid progenitors express either the ligand (Y-axis) or receptor (X-axis). **C)** Dot plot demonstrating notable ligand-receptor pairs, including co-stimulatory molecules (left) and genes involved in regulation of T cell differentiation, proliferation, or survival (right). For each interaction, MHC II^+^ erythroid progenitors express either the ligand (top, first gene listed) or receptor (bottom, second gene listed).

To further validate the potential physical interactions between erythroid progenitors and fetal T cells, we performed immunofluorescence on 22 PCW fetal liver sections. We observed direct contact between MHC II^+^ erythroid progenitors and a small portion of T cells (1.25%, Figure 6). Interestingly, areas of direct contact were often accompanied by the formation of a synapse-like structure, with polarized MHC II staining. Together, these data demonstrate that erythroid progenitors and fetal T cells have the molecular and physical potential to interact *in utero*.

**Figure 6.**
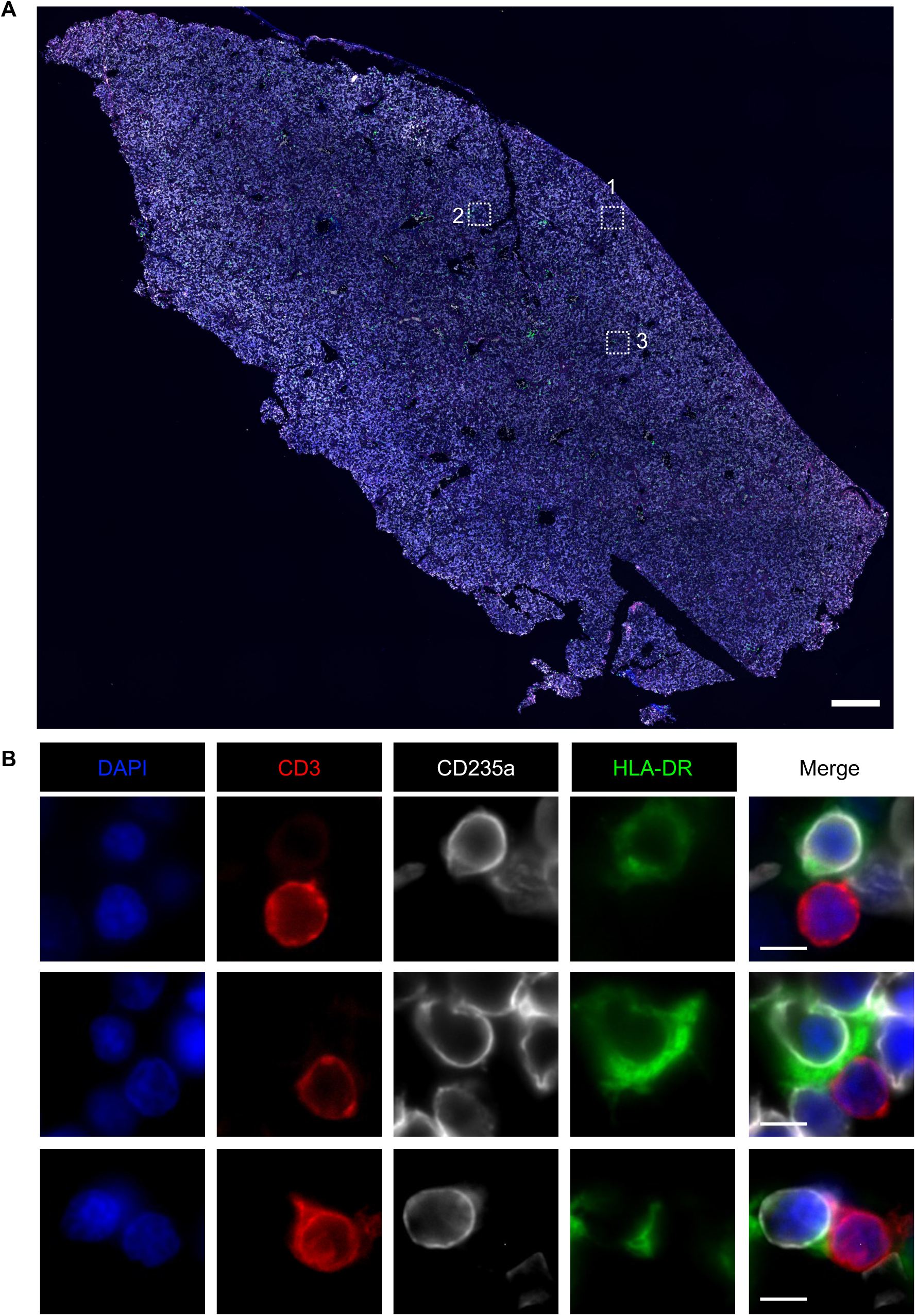
MHC II^+^ erythroid progenitors and T cells are within close physical proximity in the human fetal liver. Second trimester (22 post-conception weeks) human fetal liver was sectioned and stained for T cells (CD3), MHC II (HLA-DR), and erythroid cells (CD235a) and imaged via fluorescence microscopy. **A)** Representative 10X magnification image of one section of the fetal liver. All CD3^+^ T cells across six sections were quantified and scored based on their proximity to HLA-DR^+^ CD235a^+^ erythroid cells. Three regions of interest were selected for high magnification imaging (boxes). Scale bar = 500 µm. **B)** Representative 60X magnification images demonstrate direct contact between T cells and MHC II^+^ erythroid cells. Scale bar = 5 µm.

## Discussion

Prior functional characterizations of erythroid precursors rely on depletion via CD71 antibodies^30–32,40,42,45^. CD71 is a cell surface transferrin receptor which is expressed by a variety of cells including activated T cells^73,74^, monocytes and macrophages^75^, placental trophoblasts^32^, intestinal epithelial cells^76^, and others^77^. Our study clarifies the immunological role of erythroid progenitors while bypassing potential non-erythroid impacts of CD71-based depletion methods. Mechanistic studies have identified several candidates for immunomodulation including generation of reactive oxygen species^27,28,41,43,44^ or depletion of L-arginine via arginase-II^32,41,45^. However, many of these studies achieve only partial rescue of immune phenotypes, indicating redundant or incomplete mechanisms^28,32,45^. Our work reveals another potential mechanism by which erythroid progenitors may modulate host immunity.

Unexpectedly, we found that erythroid progenitors express MHC II antigen presentation machinery, which is typically restricted to professional antigen presenting cells. This discovery challenges the conventional view of erythroid cells as non-immunological entities and suggests a previously unrecognized role in immunomodulation. Our findings that erythroid progenitors upregulate antigen presentation machinery in response to IFN-γ underscores the capacity of these cells to modulate their activity in response to inflammatory signals.

T cell effector function is determined by the combination of costimulators and instructive cytokines expressed by antigen presenting cells. Erythroid progenitors express co-inhibitory molecules and immunoregulatory cytokines, and thus likely serve a tolerogenic role *in utero*. Given that atypical antigen presentation can mediate T_reg_ differentiation^21–24^, it is reasonable to hypothesize that erythroid progenitors contribute to the development of T_reg_ cells at the maternal-fetal interface. This hypothesis is supported by our cell-cell interaction analysis, which indicates that MHC II^+^ erythroid progenitors and fetal T_reg_ cells express complementary ligand-receptor pairs including atypical co-stimulatory molecules (Figure 5).

We observed direct physical contact between erythroid progenitors and T cells in human fetal liver samples. Our observation of synapse-like structures between MHC II^+^ erythroid progenitors and fetal T cells, along with the polarization of MHC II staining at points of contact, indicates a specialized interface for cell-cell interaction, reminiscent of the immunological synapse observed in conventional T cell interactions.

While we were somewhat surprised to find that murine erythroid cells do not express MHC II, species-specificity has been demonstrated previously in other atypical antigen presenting cells^13,14^. Mice utilize different CIITA promoters than humans, which may drive differences in MHC II expression. Furthermore, in contrast to humans, murine T cell development occurs primarily postnatally, and murine and human fetal T cells are functionally distinct^78^. This finding stresses the importance of studying antigen presentation in humans.

Our data demonstrate the potential for direct interactions between erythroid progenitors and fetal T cells. However, given the difficulty in obtaining and experimenting with human fetal tissue, it remains challenging to assess outcomes of these interactions. Future studies should focus on the functional consequences of T cell interactions to further unravel the immunobiological role of erythroid progenitors.

Broadly, these findings suggest the potential for erythroid progenitors to contribute to immune homeostasis during fetal development, and could open new avenues for understanding immune-related complications during pregnancy.

## Acknowledgments

The authors would like to thank Marisa Egan and Sunny Shin for providing THP-1 cells and protocols and advice for maintenance and activation of THP-1 cells, the Penn Cytomics and Cell Sorting facility for their help with flow cytometry and cell sorting, the Carolinas Cord Blood Bank and donors for providing umbilical cord blood, and the High Throughput Sequencing Core at the Children’s Hospital of Philadelphia for their help with bulk RNA-sequencing. This work was supported by the American Heart Association Postdoctoral Fellowship 23POST1013585 (RLC), National Institutes of Health fellowship 5T32DK007780 (RLC), National Institutes of Health IRACDA fellowship K12GM081259 (RLC), Burroughs Wellcome Fund 2022 Next Gen Pregnancy Initiative Award (KAJ), The David and Lucile Packard Foundation Fellowship for Science and Engineering (KAJ), National Institutes of Health U01 HL134696 (STC), Distinguished Chair in Pediatrics (STC), and CHOP Breakthrough Funds (STC).

## Authorship Contributions

RLC, STC, and KAJ contributed to study design and planning; HHA, MMA, STC, and LCE provided reagents, cell lines, and technical support; RLC, DS, EAK, and AC performed experiments; RLC, DS, and AC contributed to data analysis and interpretation; RLC, KRA, and TME performed secondary analysis of publicly available data; and all authors contributed to writing, editing, and reviewing of the final manuscript.

