## Supplemental Materials for "Human erythroid progenitors express antigen presentation machinery"

### Methods

#### *Flow cytometry*

Human cells: non-specific antibody binding was blocked in 10% human serum (Research Products International) and 0.5% BSA in PBS for 30 minutes on ice. Murine cells: non-specific antibody binding was blocked in 10 μg/mL anti-mouse CD16/CD32 antibody (BioXCell, clone 2.4G2) and 0.5% BSA in PBS for 30 minutes on ice. All cells were stained with antibodies for 30 minutes on ice, 1 μM Hoechst 33342 (AAT Bioquest) for 5 minutes on ice, and/or LIVE/DEAD Fixable Aqua Cell Stain Kit (Thermo Fisher) for 30 minutes on ice, washed twice in 0.5% BSA in PBS, and resuspended in 0.5% BSA in PBS. Data were acquired on LSR II (BD Biosciences) or FACSAria II (BD Biosciences) flow cytometer. Analysis was performed using FlowJo software.

#### *HUDEP-2 cell culture*

Human Umbilical cord blood-Derived Erythroid Progenitor (HUDEP-2) cells were obtained from Riken BioResource Center Cell Bank (RCB4557) through the National BioResource Project of the MEXT, Japan. Undifferentiated HUDEP-2 cells were maintained at a density of 100,000-500,000 cells/mL in StemSpan SFEM (STEMCELL) with 50 ng/mL recombinant human stem cell factor (PeproTech), $10^{-6}$ M dexamethasone (Sigma), 1 μg/mL doxycycline (Sigma), and 2 U/mL recombinant human erythropoietin (PeproTech) in a humidified, 37°C incubator at 5% $CO_2$.

#### *Human induced pluripotent stem cell culture*

Irradiated mouse embryonic fibroblasts (MEFs, R&D Systems) were seeded onto gelatin-coated plates. WT17.6 human induced pluripotent stem cell (iPSC) cultures were maintained as described previously[79] on irradiated MEFs in DMEM/F12 (Invitrogen) with 20% knockout serum replacement (Invitrogen), 2 mM L-glutamine, 1% penicillin/streptomycin, 100 μM non-essential amino acids, 100 μM beta-mercaptoethanol, and 10 ng/mL recombinant human bFGF (R&D Systems). Cultures were split weekly after incubation with TrypLe (Invitrogen) for 3 minutes and plated on fresh MEFs with 3.2 μg/mL Y27632 (ROCK inhibitor, Sigma).

#### *Orbital embryoid body differentiation*

Differentiation of iPSCs to hematopoietic progenitor cells by embryoid body differentiation was performed as previously described[79] in ultra-low attachment tissue culture plates (Corning). Embryoid bodies were cultured in StemPro 34 media (Invitrogen) supplemented with 2 mM glutamine, 50 μg/mL ascorbic acid (Sigma), 150 μg/mL transferrin (Sigma), 0.4 mM monothioglycerol (Sigma). On days 0-2, media was further supplemented with 25 ng/mL recombinant bone morphogenetic protein 4 (R&D) and 50 ng/mL VEGF (R&D). On days 2-4, media was supplemented with 25 ng/mL bone morphogenetic protein 4 (R&D), 50 ng/mL VEGF (R&D), 50 ng/mL recombinant human stem cell factor (PeproTech), 50 ng/mL recombinant human TPO (PeproTech), 50 ng/mL recombinant human Flt3 ligand (PeproTech), and 20 ng/mL recombinant human bFGF (R&D Systems). On days 4-8, media was supplemented with 50 ng/mL VEGF (R&D), 50 ng/mL recombinant human stem cell factor (PeproTech), 50 ng/mL recombinant human TPO (PeproTech), 50 ng/mL recombinant human Flt3 ligand (PeproTech), and 20 ng/mL recombinant human bFGF (R&D Systems). Hematopoietic progenitor cells were collected from embryoid body cultures 8 days post-differentiation and subsequently frozen in 40% fetal bovine serum, 10% DMSO, and 50% Iscove's Modified Dulbecco's Medium (IMDM, Gibco).

***iRBC differentiation and isolation of erythroid cells***

Frozen hematopoietic progenitor cells isolated from embryoid bodies were thawed and cultured in Iscove's Modified Dulbecco's Medium (IMDM, Gibco) with 2% human plasma (STEMCELL), 3% human AB serum (Research Products International), 1% penicillin/streptomycin, 10 µg/mL recombinant human insulin (Sigma), 200 µg/mL human holo-transferrin (Sigma), 10 ng/mL recombinant human stem cell factor (PeproTech), 1 ng/mL recombinant human IL-3 (PeproTech), 3 U/mL recombinant human erythropoietin (PeproTech), and 3 U/mL heparin (Sigma). The next day, non-adherent cells were collected centrifuged at 300 *x g* for 5 minutes,. Media was removed and replaced with IMDM with 2% human plasma, 3% human AB serum, 1% penicillin/streptomycin, 10 µg/mL recombinant human insulin, 1 mg/mL human holo-transferrin, 10 ng/mL recombinant human stem cell factor, 3 U/mL recombinant human erythropoietin, and 3 U/mL heparin. 3 days later, non-adherent cells were collected and centrifuged at 300 *x g* for 5 minutes. Pellet was resuspended in 0.5% bovine serum albumin (BSA) in PBS with anti-CD235a magnetic microbeads (Miltenyi Biotec) for 15 minutes at 4°C. Erythroid cells were isolated using LS magnetic columns (Miltenyi Biotec) following manufacturer’s instructions.

***TGF-β1 secretion***

TGF-β1 levels in HUDEP-2 culture supernatants or non-conditioned media were measured using the human TGF-β1 Quantikine ELISA kit (R & D Systems) according to the manufacturer's instructions using a SpectraMAX 190 plate reader (Molecular Devices).

***THP-1 cell culture and stimulation***

The human acute monocytic leukemia cell line THP-1 was obtained from Sunny Shin (University of Pennsylvania). THP-1 cells were maintained at a density of 300,000-900,000 cells/mL in RPMI 1640 (Gibco) with 10% heat-inactivated fetal bovine serum (HyClone) and 0.05 mM β-mercaptoethanol (Gibco) in a non-tissue culture-treated dish in a humidified, 37°C incubator at 5% $CO_2$. THP-1 cells were differentiated into adherent macrophage-like cells by culturing 700,000 cells/ml in RPMI 1640 with 10% heat-inactivated fetal bovine serum and 200 nM phorbol myristate acetate (InvivoGen) for 24 hours in a tissue culture-treated dish. For flow cytometry experiments, adherent macrophage-like cells were removed from the dish with a cell scraper prior to staining.

***Cathepsin protease activity assay***

Cathepsin activity was assessed using the SensoLyte 520 Cathepsin S and D Assay Kits (Anaspec). For each technical replicate for each cathepsin enzyme, 500,000 undifferentiated, unstimulated HUDEP-2 or THP-1 cells were lysed and cathepsin activity was assessed according to the manufacturer’s instructions. Fluorescence was measured after 30 minutes using a SpectraMAX 190 plate reader (Molecular Devices). Relative fluorescence units were obtained by subtracting baseline fluorescence from the obtained readings.

***Antigen internalization assays***

100,000 HUDEP-2 cells from 0 or 6 days post-differentiation were incubated with 10 µg/mL ovalbumin-Alexa Fluor 488 (Invitrogen) at 37°C for 1.5 hr. Cells were washed three times in 0.5% bovine serum albumin in PBS. Fluorescence was measured via flow cytometry.

***Mice***

Wild type C57BL/6 mice (Jackson Laboratories) were bred and maintained at the University of Pennsylvania. Mice were euthanized with $CO_2$ in compliance with the University of Pennsylvania Institutional Animal Care and Use Committee protocols. Bone marrow was isolated as described previously[80]. Briefly, femur and tibia from 6-8 week female C57BL/6 mice were centrifuged in nested microcentrifuge tubes at 10,000 *x g* for 15 seconds at 4°C, then bone marrow was resuspended in 0.5% bovine serum albumin (BSA) in PBS. Spleens were dissected from neonatal mice (postnatal day 5) and homogenized through a 70 µM cell strainer (Celltreat). Splenocytes were centrifuged at 300 *x g* for 5 minutes at 4°C, resuspended in RPMI 1640 (Gibco) with 10% heat-inactivated fetal bovine serum (HyClone), and plated at 5 x $10^5$ cells/mL. Splenocytes were stimulated with 200 ng/mL recombinant mouse interferon-gamma (BD Biosciences) for 24 hours in a humidified, 37°C incubator at 5% $CO_2$.

***CIITA promoter-specific qPCR analysis***

1x$10^6$ undifferentiated HUDEP-2 cells were stimulated with 50 ng/mL recombinant human interferon-gamma (PeproTech) for 6 hours. Stimulated cells and unstimulated controls were resuspended in TRIzol (Invitrogen) and RNA was isolated via phase separation using 1-bromo-3-chloropropane. Aqueous layer was precipitated in isopropanol and RNA was washed in 70% ice-cold ethanol, then resuspended in nuclease-free water. cDNA synthesis was completed with iScript Reverse Transcriptase (Bio-Rad) using 1 ug of RNA per reaction in a ProFlex PCR System (Thermo Fisher). qPCR primers were designed with a forward primer specific to the unique first exon adjacent to each promoter and a shared reverse primer specific to exon 2. qPCR was performed using SYBR Green (Thermo Fisher) in a QuantStudio 3 Real-Time PCR System (Thermo Fisher). Reactions were performed in technical duplicates for stimulated cells, unstimulated control, and a water-only control. GAPDH was used as a housekeeping gene.

***In-vitro erythropoiesis in HUDEP-2 cells***

HUDEP-2 cells were centrifuged at 200 x *g* for 5 minutes and resuspended at a density of 500,000 cells/mL in Iscove's Modified Dulbecco's Medium (IMDM, Gibco) with 2% human plasma (STEMCELL), 3% human AB serum (Research Products International), 10 µg/mL recombinant human insulin (Sigma), 200 µg/mL human holo-transferrin (Sigma), 10 ng/mL recombinant human stem cell factor (PeproTech), 1 ng/mL recombinant human IL-3 (PeproTech), 2 U/mL recombinant human erythropoietin (PeproTech), and 1 µg/mL doxycycline (Sigma). Every 3 days, cells were counted and brought to a density of 500,000 cells/mL by adding IMDM with 2% human plasma (STEMCELL), 3% human AB serum (Research Products International), 10 µg/mL recombinant human insulin (Sigma), 1000 µg/mL human holo-transferrin (Sigma), and 3 U/mL recombinant human erythropoietin (PeproTech). Insulin, holo-transferrin, and erythropoietin concentrations were calculated proportional to the total volume, while plasma and serum were added proportional to the additional volume added.

***Immunofluorescence***

Study subjects provided written informed consent to participate in this study. The institutional review boards at the Children's Hospital of Philadelphia and the University of Pennsylvania approved the study protocol. Second trimester (22 post-conception weeks) human fetal liver was obtained following informed consent and incubated in 4% paraformaldehyde for 24 hours at 4°C. Sample was sequentially immersed in 10%, 20%, and 30% sucrose until sunk, then then embedded in OCT (Sakura) and cryosectioned (10 µm). Sections were washed twice in PBS, then three times in PBS containing 0.4% Triton X-100 (PBST) for 10 minutes each, then blocked with PBS containing 0.4% Triton X-100, 1% BSA, and 2% normal goat serum (blocking solution) for 1 hour. Samples were incubated with primary antibodies overnight at 4°C (Supplemental Table 1) 1:100 in blocking solution. Samples were washed three times in PBST for 10 minutes each, then incubated with secondary antibodies (1:500) and DAPI for 2 hours at room temperature.

**Supplementary Table 1. Flow cytometry and immunofluorescence antibodies**

| Target | Clone | Vendor |
|---|---|---|
| CD235a (human) | HIR2 (FC) or<br>JC159 (IF) | BioLegend or<br>Cell Signaling Technology |
| CD71 (human) | OKT-9 | eBioscience |
| CD45 (human) | 2D1 | BioLegend |
| CD34 (human) | 581 | BioLegend |
| HLA-DR (human) | LN3 | BioLegend |
| HVEM (human) | 122 | BioLegend |
| CD80 (human) | 2D10 | BioLegend |
| CD86 (human) | IT2.2 | BioLegend |
| ICOSL (human) | 2D3 | BioLegend |
| PD-L1 (human) | 29E.2A3 | BioLegend |
| VISTA (human) | B7H5DS8 | eBioscience |
| Galectin-9 (human) | 9M1-3 | BioLegend |
| CD3 (human) | OKT3 or UCHT1 | BioLegend |
| CD4 (human) | OKT4 | BioLegend |
| CD69 (human) | FN50 | BioLegend |
| PD-1 (human) | EH12.2H7 | BioLegend |
| Ter119 (mouse) | TER-119 | BioLegend |
| CD71 (mouse) | RI7217 | BioLegend |
| I-A/I-E (mouse) | 2G9 | BD Biosciences |
| CD34 (mouse) | HM34 | BioLegend |
| anti-mouse IgG2b AF488 | N/A | BioLegend |
| anti-mouse IgG1 AF594 | N/A | BioLegend |
| anti-rabbit Cy5 | N/A | Jackson ImmunoResearch |

A

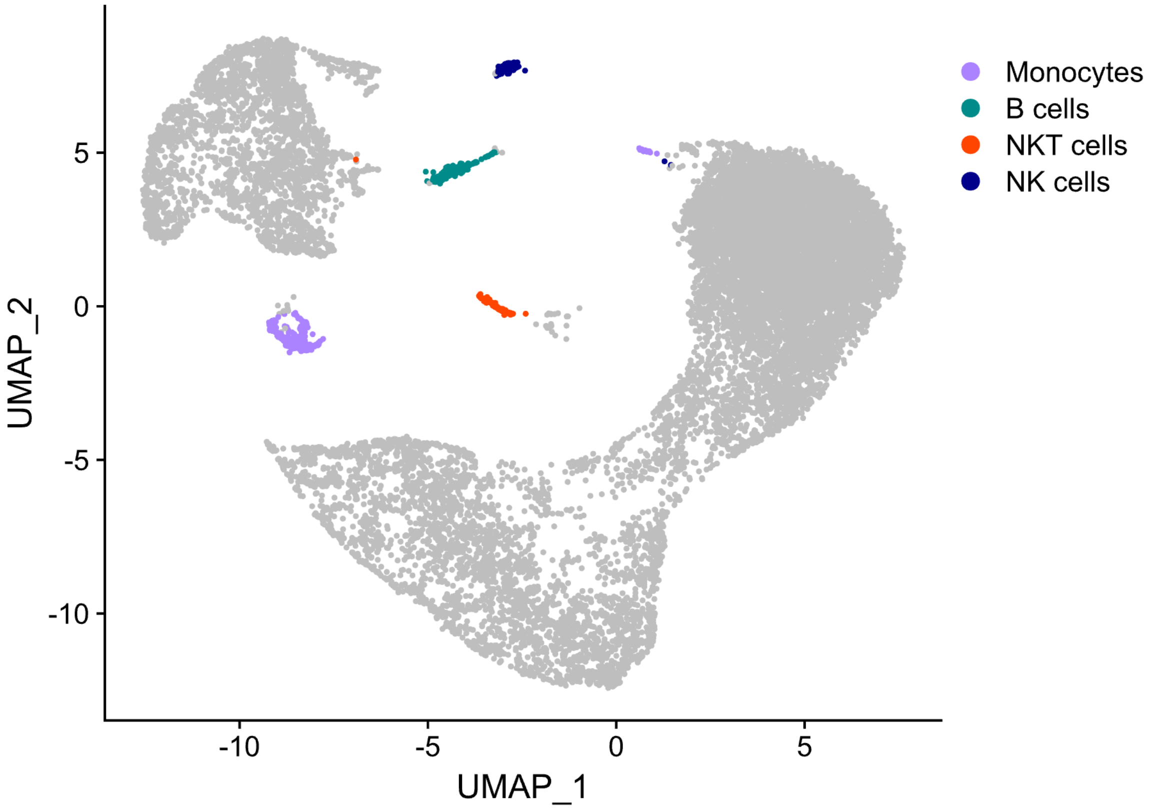


B

| Cell Type | Markers |
|---|---|
| *CD4 T cells* | CD3 gamma chain (ENSG00000160654), CD3 delta chain (ENSG00000167286), CD3 epsilon chain (ENSG00000198851), CD4 (ENSG00000010610) |
| *CD8 T cells* | CD3 gamma chain (ENSG00000160654), CD3 delta chain (ENSG00000167286), CD3 epsilon chain (ENSG00000198851), CD8a (ENSG00000153563), CD8b (ENSG00000172116) |
| **NKT cells** | CD3 gamma chain (ENSG00000160654), CD3 delta chain (ENSG00000167286), CD3 epsilon chain (ENSG00000198851), CD28 (ENSG00000178562), CD161 (ENSG00000111796) |
| **B cells** | CD19 (ENSG00000177455), CD20 (ENSG00000156738), HLA-DRA (ENSG00000204287) |
| *Plasma cells* | CD19 (ENSG00000177455), CD27 (ENSG00000139193), CD38 (ENSG00000004468) |
| **Monocytes** | CD11c (ENSG00000140678), CD14 (ENSG00000170458), HLA-DRA (ENSG00000204287) |
| **NK cells** | CD45 (ENSG00000081237), CD56 (ENSG00000149294), CD94 (ENSG00000134539) |
| *Basophils* | CD38 (ENSG00000004468), CD123 (ENSG00000185291), CD294 (ENSG00000183134) |
| *Neutrophils* | CD16a (ENSG00000203747), CD16b (ENSG00000162747), CD66b (ENSG00000124469) |
| *Eosinophils* | CD66b (ENSG00000124469), CD294 (ENSG00000183134) |
| *Dendritic cells* | CD11c (ENSG00000140678), CD38 (ENSG00000004468), CD123 (ENSG00000185291), HLA-DRA (ENSG00000204287) |

**Supplemental Figure 1. Exclusion of non-erythroid cells from single cell RNA-sequencing analysis. A)** UMAP plot of single-cell RNA-sequencing data of primary human umbilical cord cells. Data from Huang, *et al*. (GSE150774). Clusters corresponding to doublets with erythroid cells. **B)** Markers chosen to assess the presence of each immune cell type. Cell types in bold were found contaminating the sequencing data and corresponding clusters were removed from subsequent analysis.

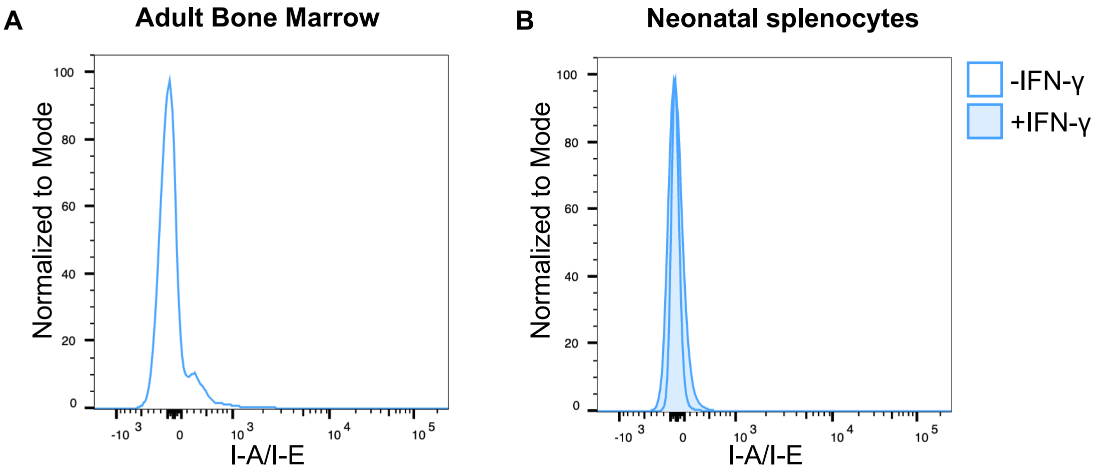


**Supplemental Figure 2. Murine erythroid cells do not express MHC class II molecules.** Flow cytometry of **(A)** adult bone marrow or **(B)** neonatal splenocytes demonstrates lack of I-A/I-E expression on murine erythroid cells (gated on live Ter119$^{+}$ cells). Neonatal splenocytes were stimulated with 200 ng/mL interferon-γ (IFN-γ) for 24 h.

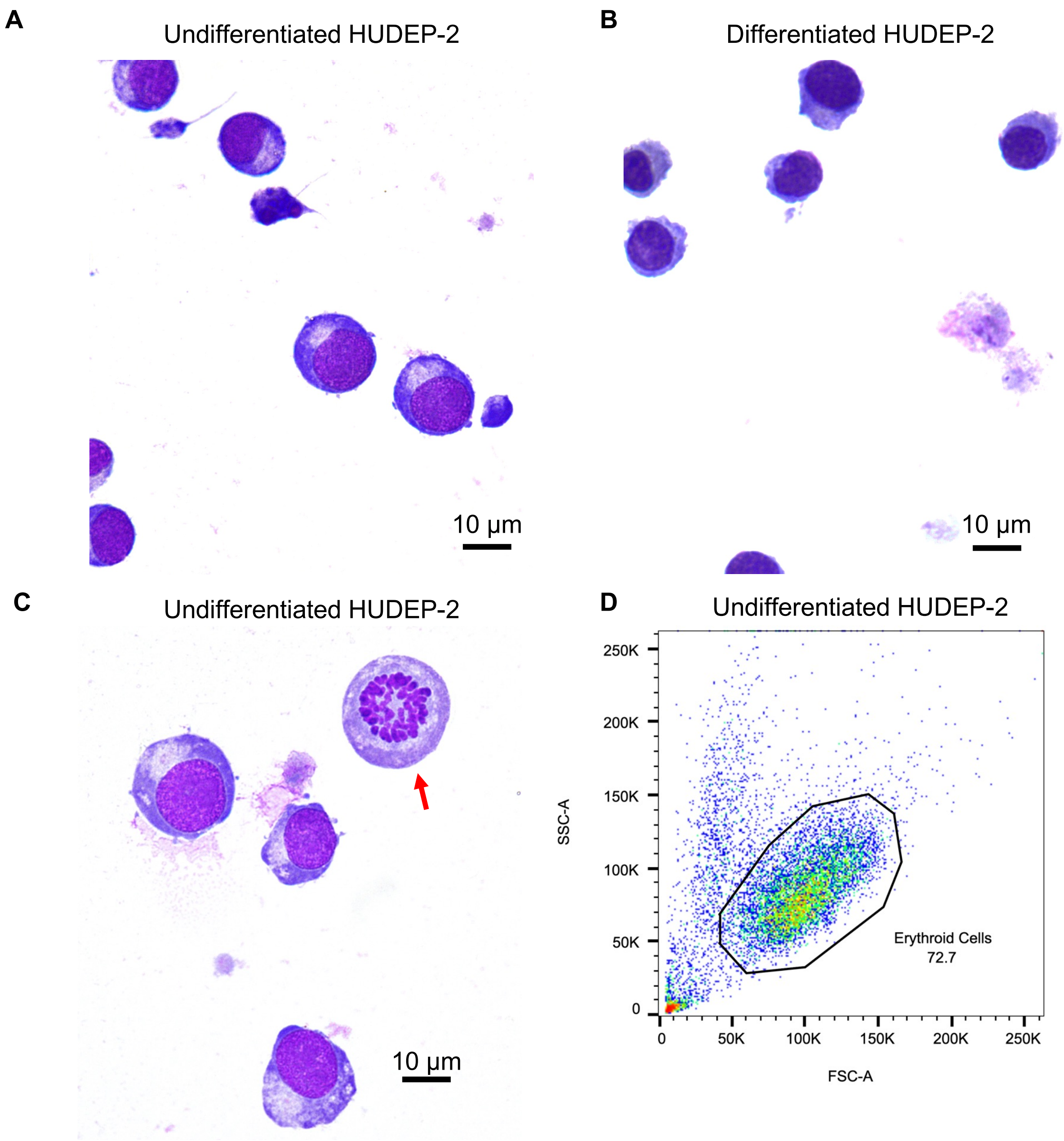


**Supplemental Figure 3. HUDEP-2 differentiation.** Hemacolor-stained smears of **A)** undifferentiated HUDEP-2 cells or **B)** HUDEP-2 cells six days post-differentiation. Scale bars = 10 µm. **C)** Hemacolor-stained smears of undifferentiated HUDEP-2 cells reveal a mixture of early erythroid cells as well as rare myeloid-like cells (red arrow). Scale bar = 10 µm. **D)** Flow cytometry of undifferentiated HUDEP-2 cells. Non-granular erythroid cells are selected via forward and side scatter as depicted to exclude rare myeloid-like cells as seen in (C).

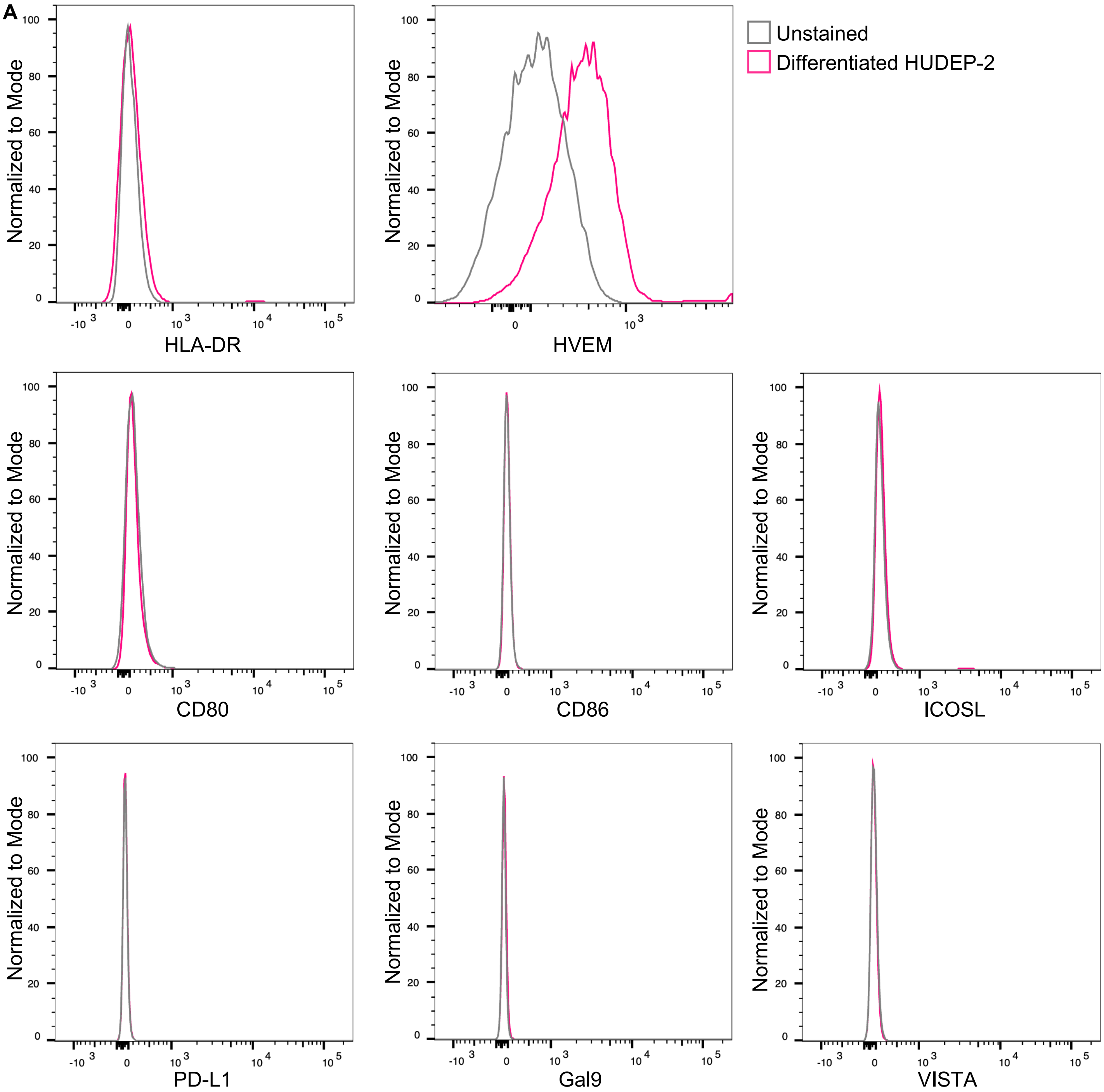


**Supplemental Figure 4. Erythroid precursors do not express MHC class II machinery or co-stimulatory markers. A)** Flow cytometry of HUDEP-2 cells six days post-differentiation reveals that erythroid precursors do not express MHC class II (HLA-DR) or co-stimulatory markers, except for HVEM. n = 3 technical replicates.

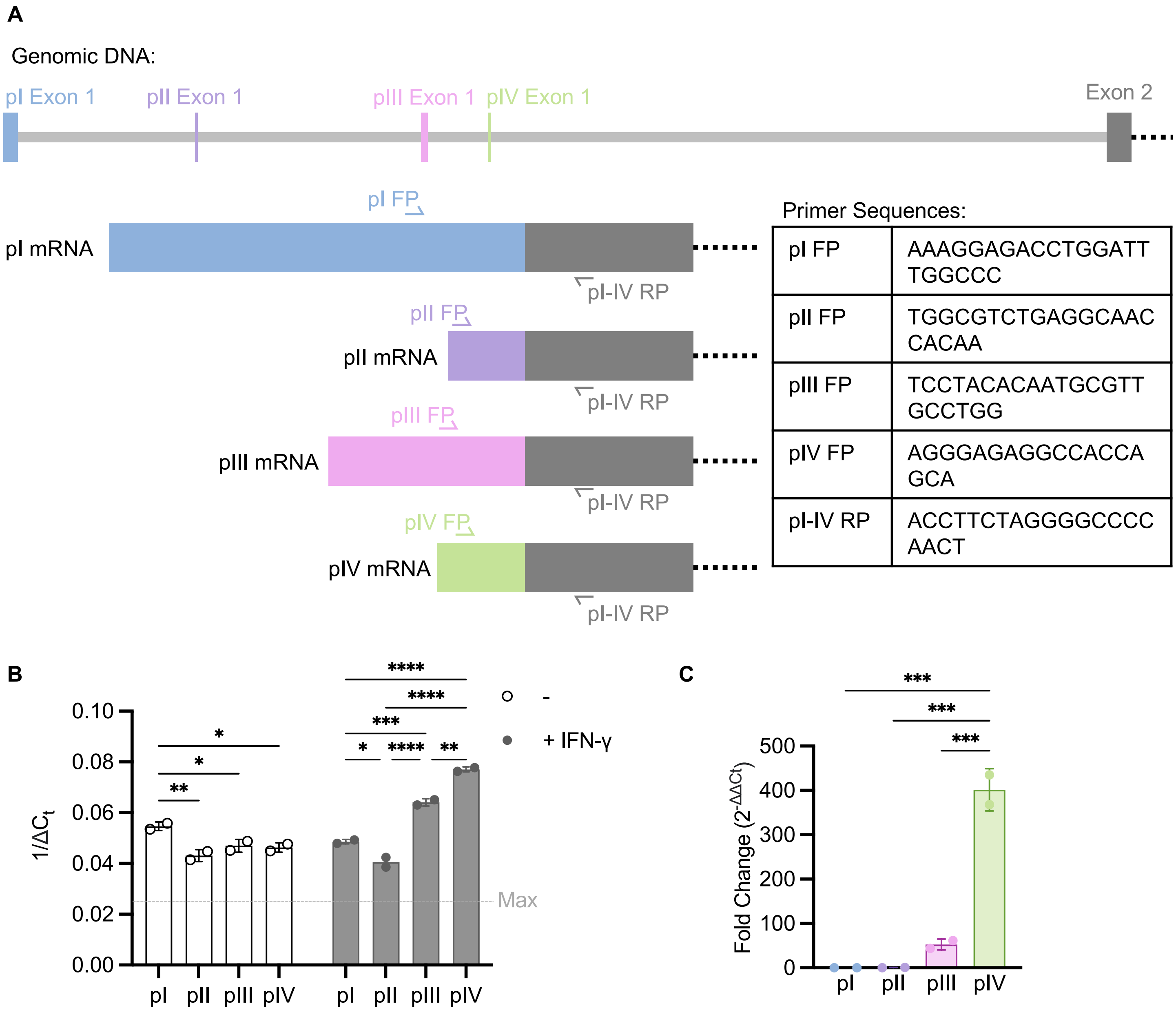


**Supplemental Figure 5. Erythroid progenitors drive baseline expression of CIITA via promoter I.** qPCR of undifferentiated HUDEP-2 cells stimulated with or without interferon-gamma (IFN-γ, 50 ng/mL, 6h) demonstrates that baseline expression of CIITA is driven primarily via pI, while upregulation of CIITA after IFN-γ stimulation is driven by pIV and pIII. **A)** Schematic of primer design. Each promoter I-IV (pI-pIV) has a unique forward primer in first exon and a common reverse primer in second exon. **B)** Cycle threshold ($C_t$) compared to GAPDH for each condition. Dashed line represents maximum number of cycles (40). Two-way ANOVA. **C)** Fold change of IFN-γ-stimulated condition compared to unstimulated control. One-way ANOVA. *P<0.0332, **P < 0.002, ***P < 0.0002, ****P < 0.0001. n = 2 technical replicates, representative of two independent experiments.

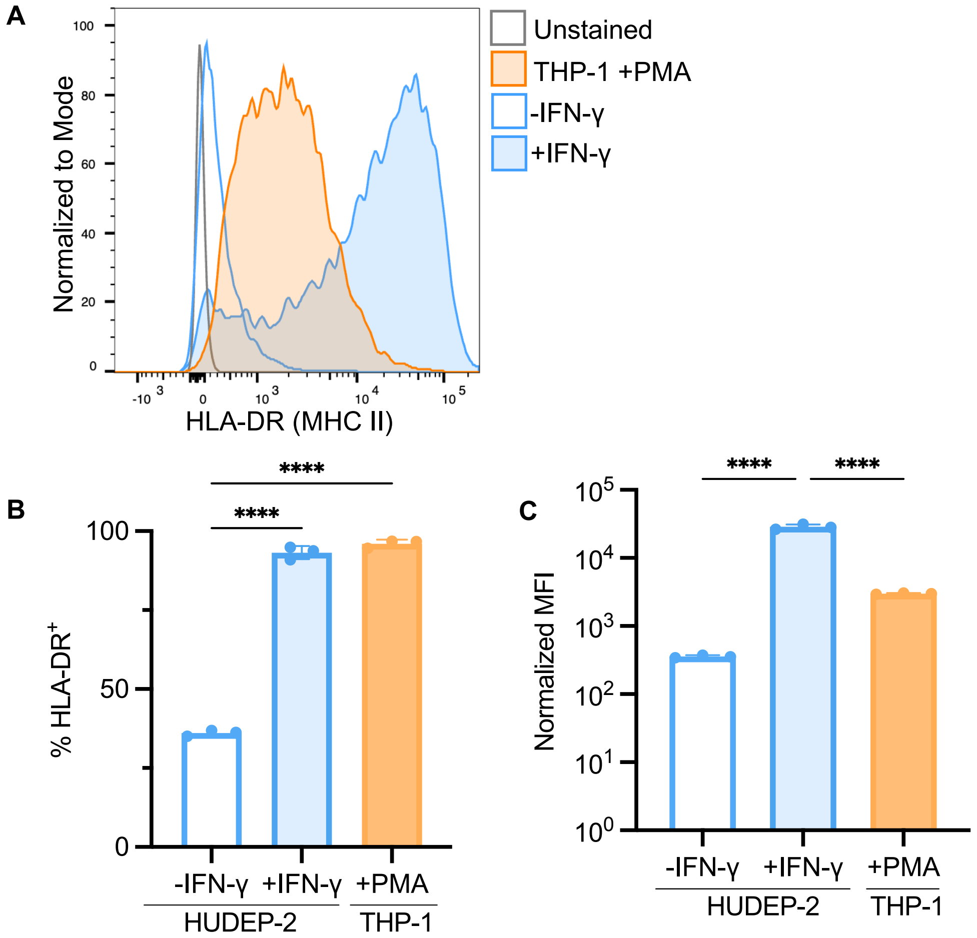


**Supplemental Figure 6. Erythroid progenitors express lower levels of HLA-DR than professional antigen presenting cells. A)** Flow cytometry of undifferentiated, unstimulated HUDEP-2 cells or phorbol myristate acetate (PMA) activated THP-1 cells reveals that erythroid progenitors express lower surface levels of MHC class II (HLA-DR) than activated antigen presenting cells at baseline. After stimulation with interferon-gamma (IFN-γ, 50 ng/mL, 72 hr), surface expression of HLA-DR in HUDEP-2 cells increases to levels higher than activated THP-1. **B)** Quantification of (A). Graph indicates percentage of HLA-DR-positive cells. n = 3 technical replicates. Mean ± SD. One-way ANOVA. ****$P < 0.0001$. **C)** Quantification of (A). Graph indicates mean fluorescence intensity. Mean ± SD. One-way ANOVA. ****$P < 0.0001$. n = 3 technical replicates.

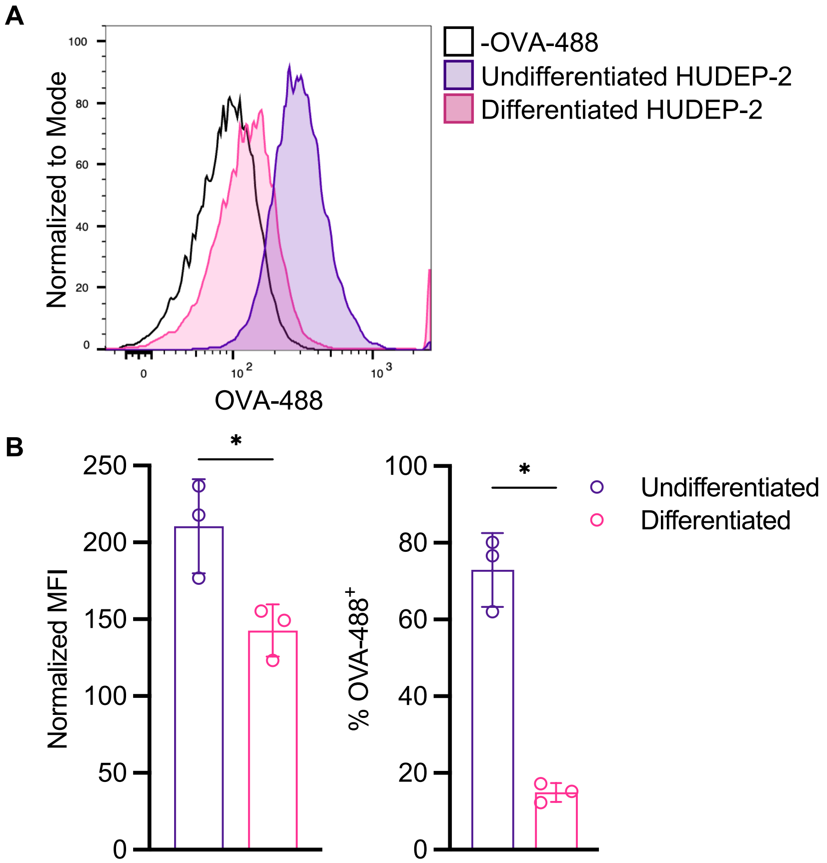


**Supplemental Figure 7. Erythroid progenitors internalize antigen. A)** Undifferentiated HUDEP-2 or differentiated HUDEP-2 (six days post-differentiation) were incubated with 10 µg/mL ovalbumin-Alexa Fluor 488 (OVA-488) at 37°C for 1.5 hr. Fluorescence measured via flow cytometry. **B)** Quantification of (A). Graphs indicate mean fluorescence intensity normalized to –OVA-488 controls or percentage of OVA-488$^+$ cells. Mean ± SD. One-way ANOVA. *P < 0.03. n = 3 technical replicates.

**A**

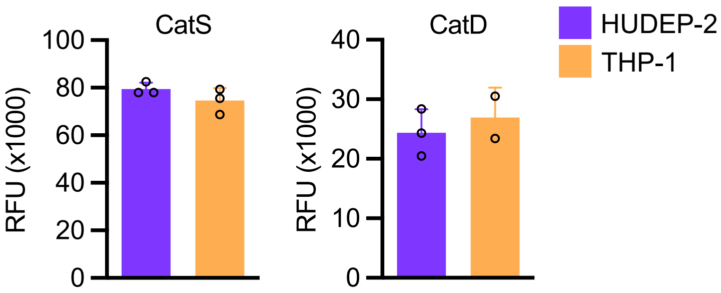


**Supplemental Figure 8. Human erythroid cells possess active endo-lysosomal proteases. A)** Cathepsin S and cathepsin D activity was assessed in undifferentiated, unstimulated HUDEP-2 cells and THP-1 cells using fluorometric cathepsin substrates. Background fluorescence was subtracted to obtain relative fluorescence values. Mean ± SD. Unpaired, two-tailed t-test. n = 3 technical replicates per experiment, representative of 3 independent experiments.

**A**

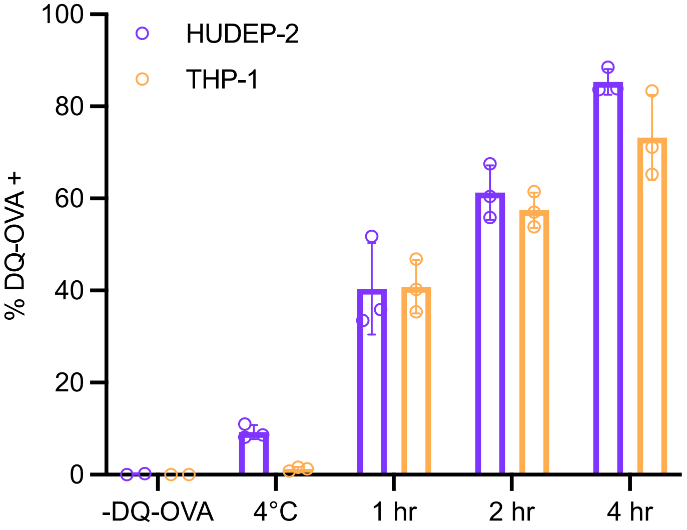


**Supplemental Figure 9. Erythroid progenitors process antigens**. **A)** Undifferentiated, unstimulated HUDEP-2 cells or THP-1 cells were incubated with 10 µg/mL DQ-OVA at 4°C for 4 hr or 37°C for indicated times. Graph indicates percentage of DQ-OVA-positive cells. Mean ± SD. Two-way ANOVA. n = 3 technical replicates per experiment, representative of 3 independent experiments.

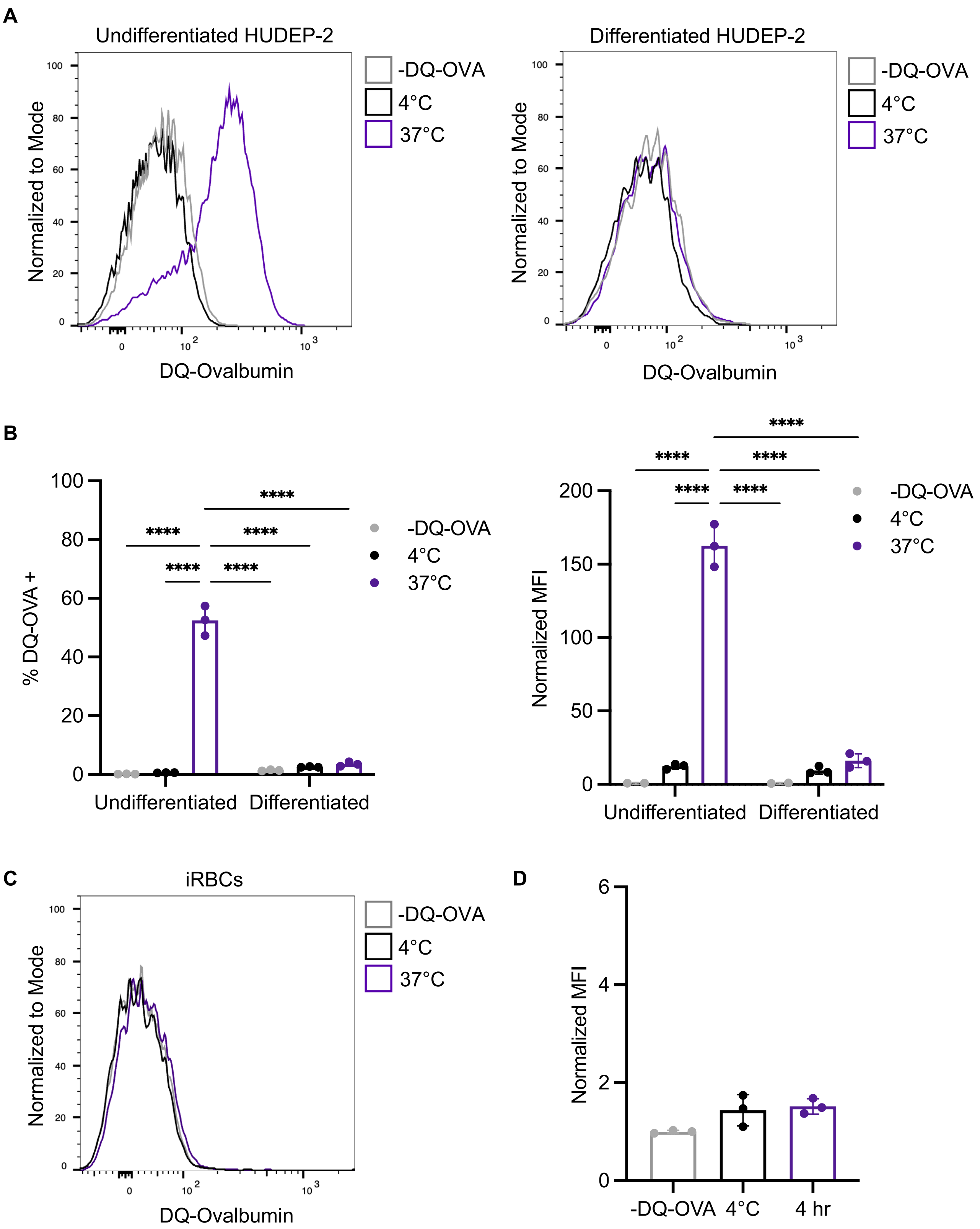


**Supplemental Figure 10. Erythroid precursors do not process antigens. A**) Undifferentiated HUDEP-2 or differentiated HUDEP-2 (six days post-differentiation) were incubated with 10 µg/mL direct quenched-ovalbumin (DQ-OVA) at 37°C or 4 °C for 4 hr. Fluorescence measured via flow cytometry indicates relief of self-quenching & successful receptor-mediated endocytosis/proteolytic digestion. **B)** Quantification of (A). Graphs indicate mean fluorescence intensity normalized to -DQ-OVA control or percentage of DQ-OVA$^+$ cells. Mean ± SD. One-way ANOVA. ****P < 0.0001. n = 3 technical replicates. **C)** iRBCs were incubated with 10 µg/mL DQ-OVA at 37°C or 4 °C for 4 hr. **D)** Quantification of (C). Graph indicates mean fluorescence intensity normalized to -DQ-OVA control. Mean ± SD. One-way ANOVA. n = 3 technical replicates.

**A**

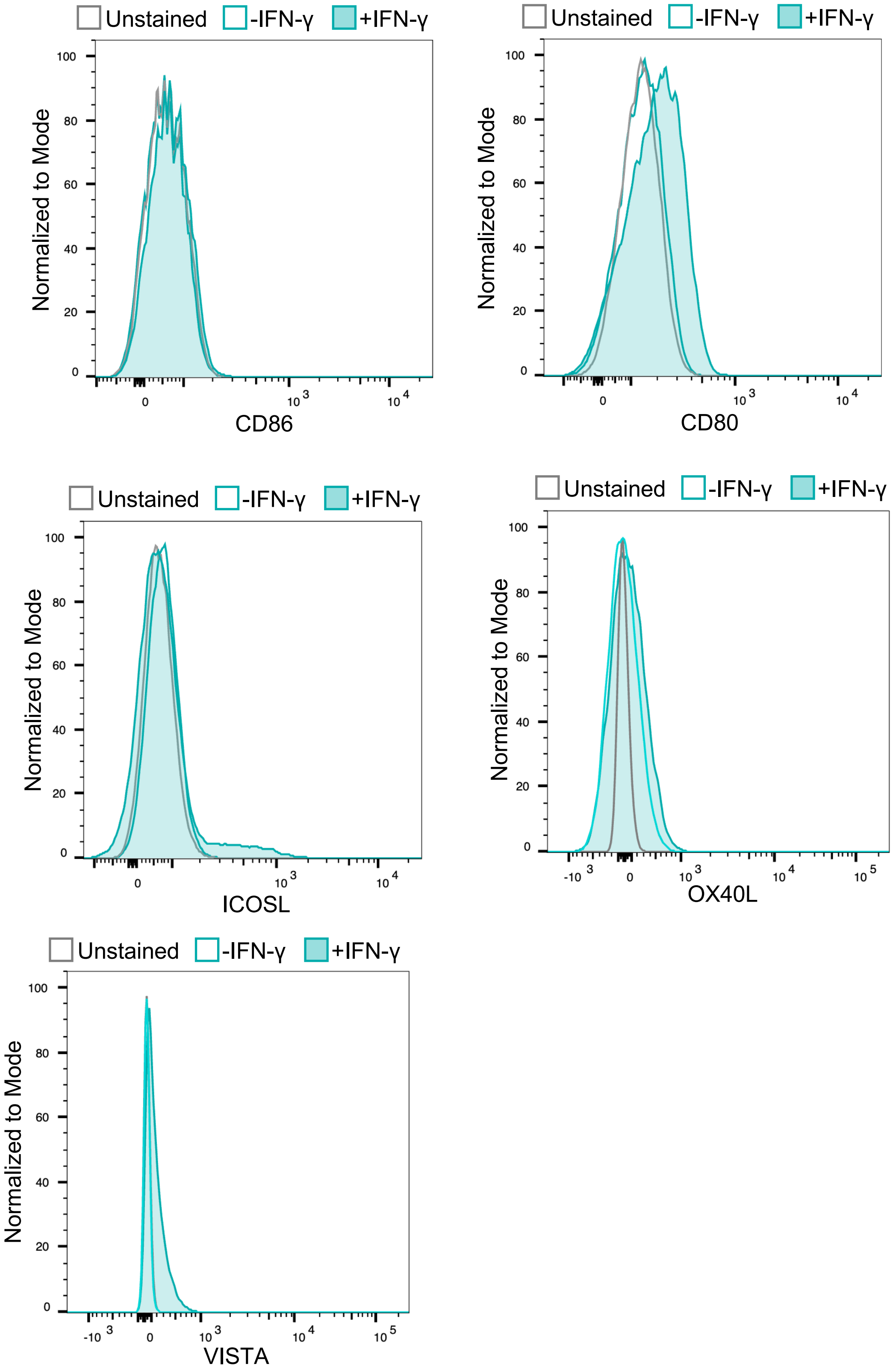


**Supplemental Figure 11. Erythroid progenitors do not express classical co-stimulators. A)** Flow cytometry of undifferentiated HUDEP-2 cells reveals that erythroid progenitors do not express classical co-stimulatory molecules (ICOSL, CD80, and CD86) or other co-stimulatory molecules (VITSA and OX40L), even upon stimulation with interferon-gamma (IFN-γ, 50 ng/mL, 72h). Representative flow plots from 3 replicates.

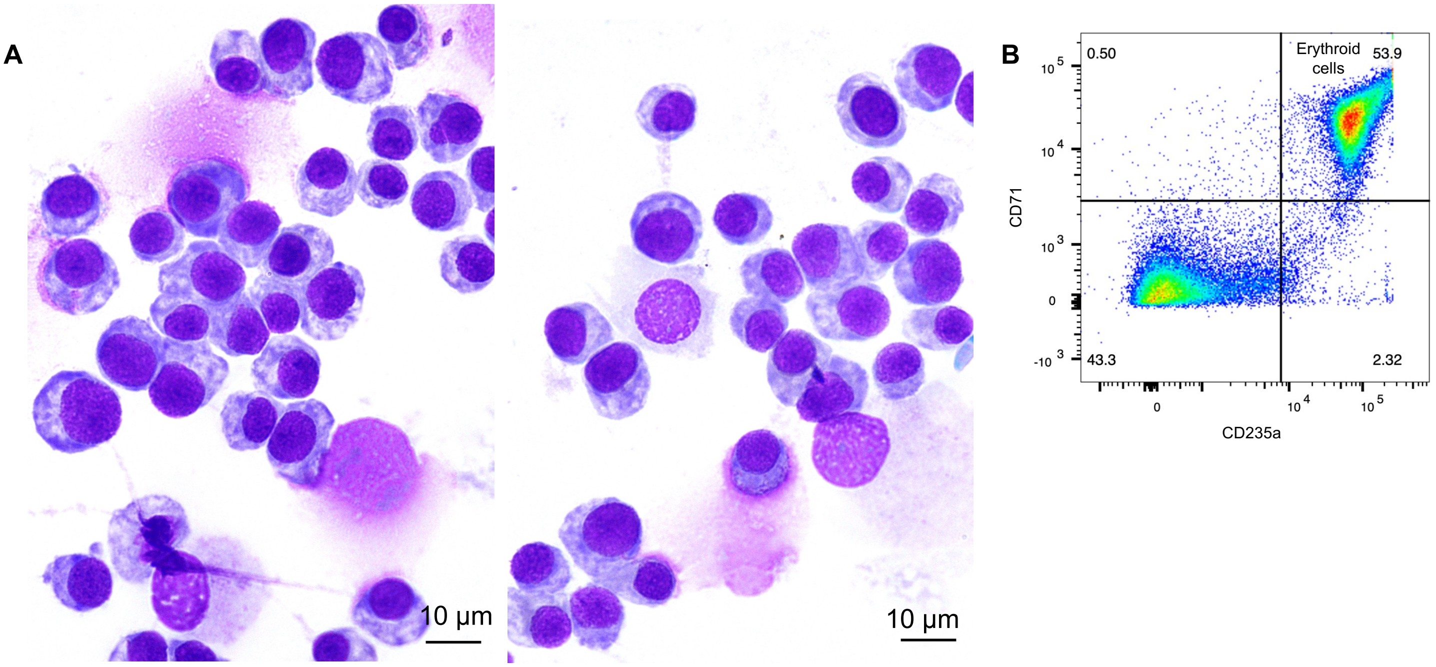


**Supplemental Figure 12. Induced pluripotent stem cell derived erythroid progenitors. A)** Hemacolor-stained smears of iRBCs 3 days post-differentiation reveal a mixture of early-to-mid erythroid cells as well as stem-like cells. Scale bars = 10 µm. **d)** Flow cytometry of iRBCs 3 days post-induction of erythropoiesis. Erythroid cells are selected via CD235a expression and/or magnetic-activated cell sorting.

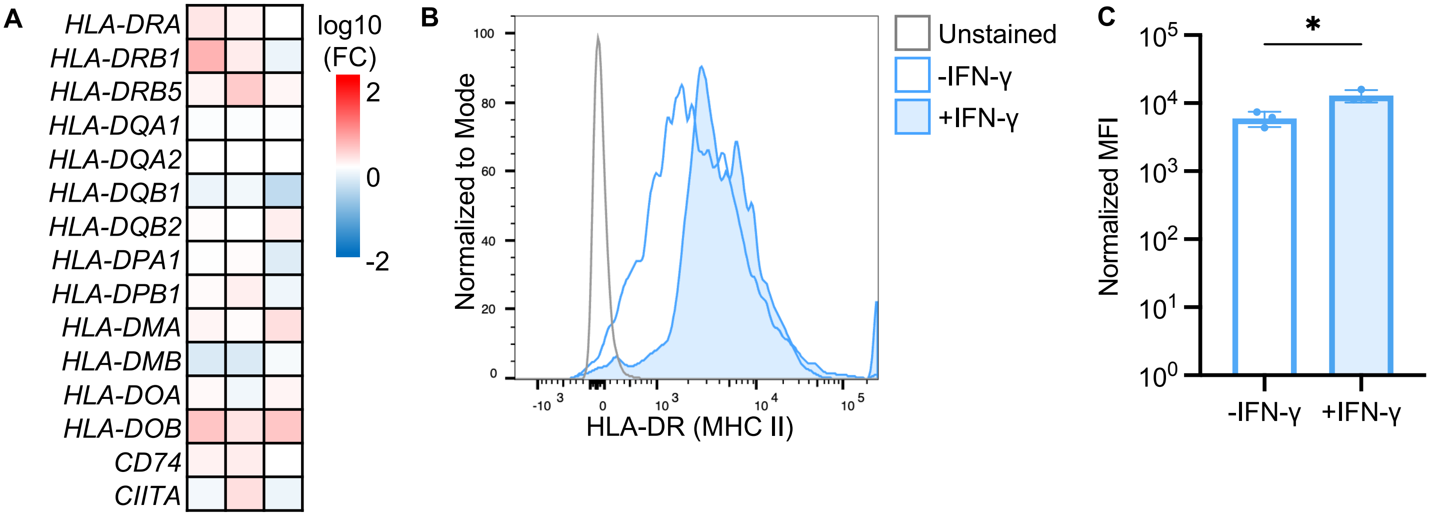


**Supplemental Figure 13. iRBCs express MHC class II antigen presentation machinery but are less responsive to interferon stimulation. A)** Bulk RNA-sequencing of iRBCs 3 days post-differentiation reveals that iRBCs do not upregulate expression of MHC class II molecules and associated machinery in response to stimulation with interferon-gamma (IFN-γ, 50 ng/mL, 6h). Heatmap displays fold-change (FC) compared to unstimulated control for three replicates. **B)** Flow cytometry of iRBCs reveals that erythroid progenitors slightly upregulate surface expression of MHC class II (HLA-DR) in response to stimulation with interferon-gamma (IFN-γ, 50 ng/mL, 72 h). **C)** Quantification of (B). Graph indicates mean fluorescence intensity normalized to unstained control. Mean ± SD. Unpaired, two-tailed t-test. *P<0.03. n = 3 technical replicates.

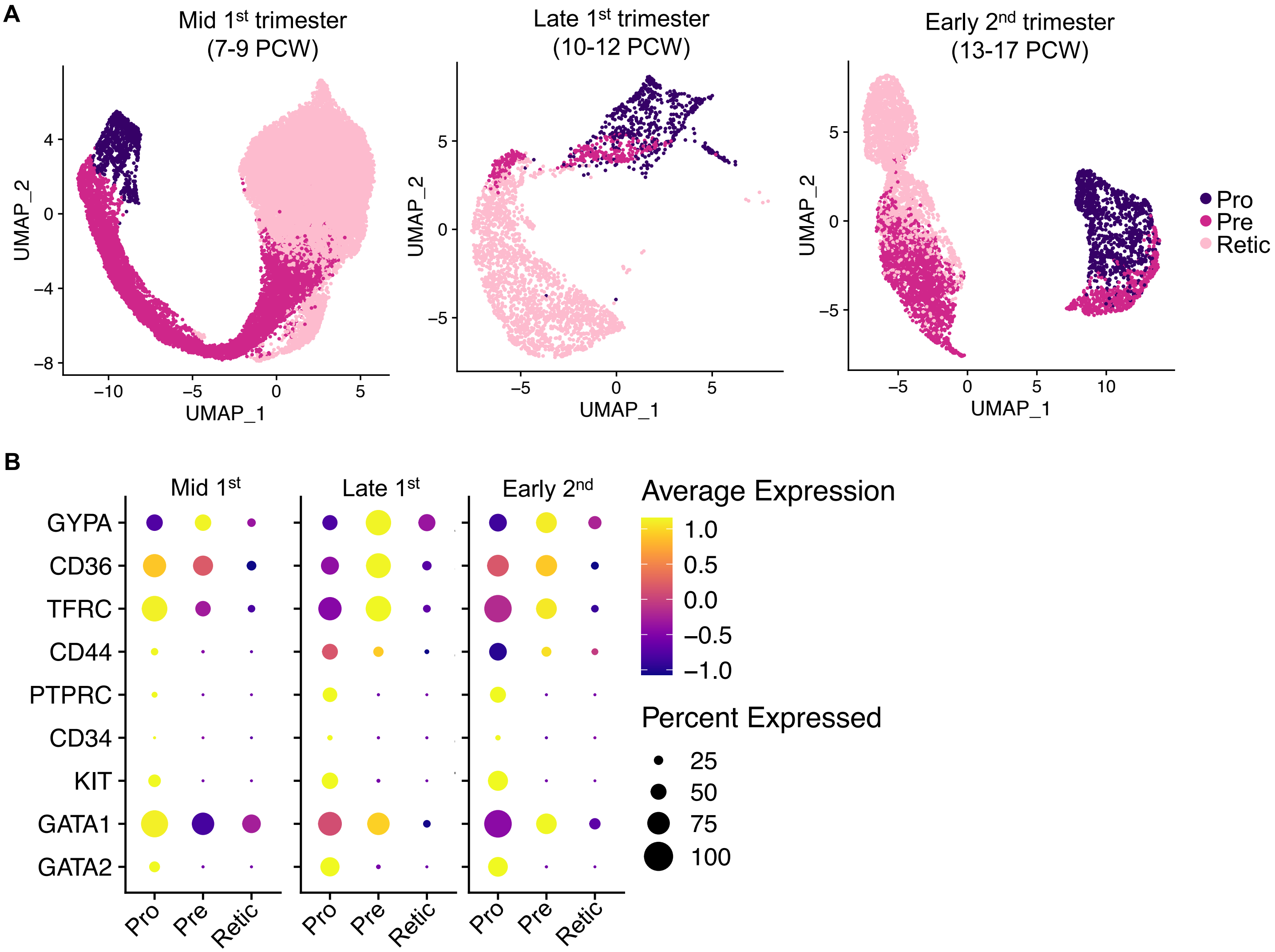


**Supplemental Figure 14. Analysis of erythroid progenitors throughout pregnancy. A)** UMAP plot of single-cell RNA-sequencing data of primary human fetal erythroid cells. Data from Suo, *et al*. Data was separated into mid first trimester samples (7-9 post-conception weeks (PCW)), late first trimester samples (10-12 PCW), or early second trimester samples (13-17 PCW) and filtered on erythroid cells. Clusters were divided erythroid progenitors (pro), precursors (pre), or reticulocytes (retic). **B)** Analysis parameters and clustering of single-cell RNA-sequencing data was validated with known surface markers of erythropoiesis and transcription factors differentially expressed during RBC development. *GYPA* = CD235a, *TRFC* = CD71, *PTPRC* = CD45, *KIT* = CD117.

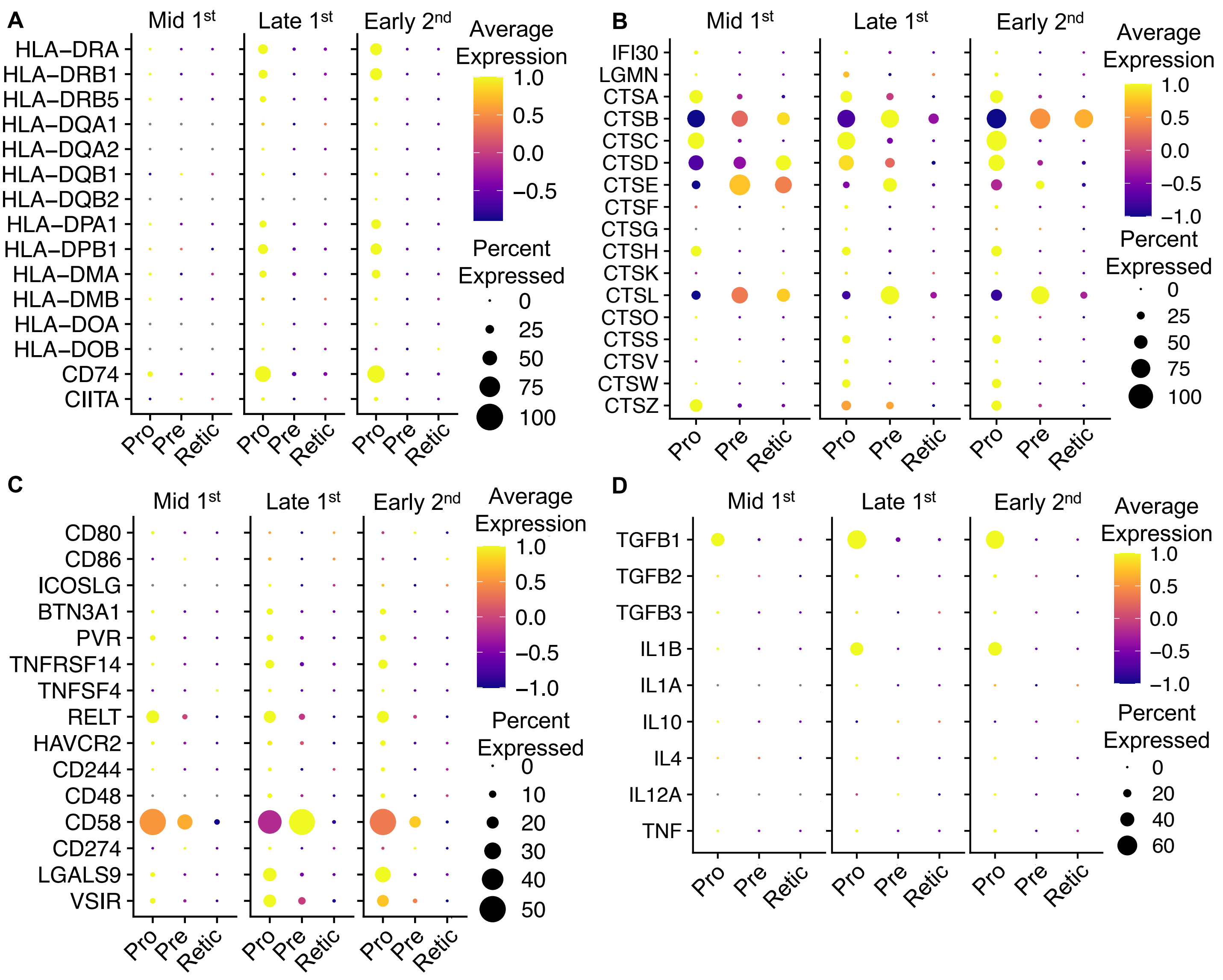


**Supplemental Figure 15. Antigen processing and presentation machinery throughout human fetal development.** Single cell RNA-sequencing of primary fetal erythroid cells reveals expression of **A)** MHC class II and associated machinery, **B)** endo-lysosomal proteases, **C)** co-stimulatory markers, and **D)** instructive cytokines in erythroid progenitors. Data from Suo, *et al*. Data was separated into mid first trimester samples (7-9 post-conception weeks (PCW)), late first trimester samples (10-12 PCW), or early second trimester samples (13-17 PCW) and filtered on erythroid cells. Clusters were divided erythroid progenitors (pro), precursors (pre), or reticulocytes (retic).

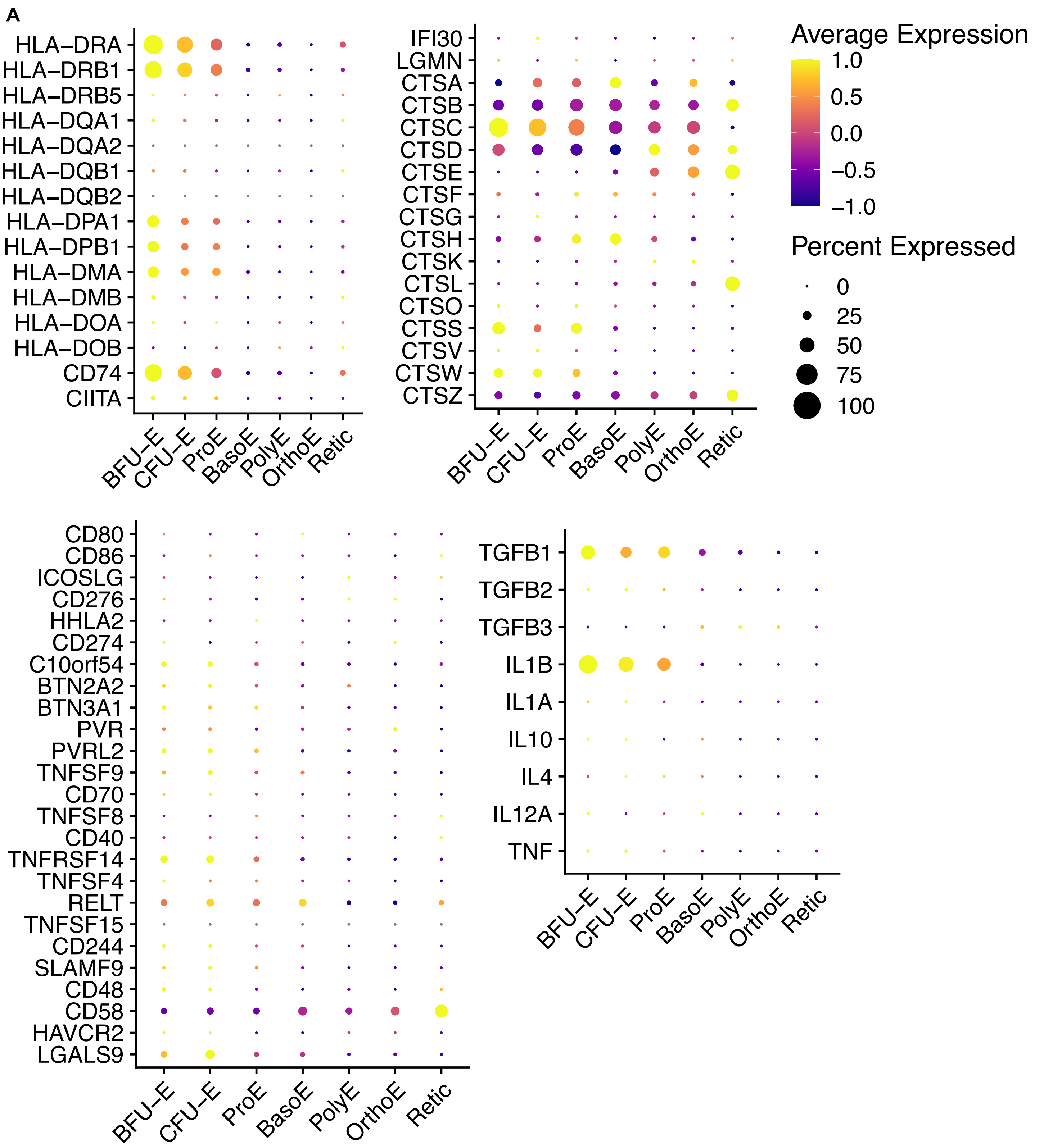


**Supplemental Figure 16. Human fetal liver erythroid progenitors express MHC class II antigen processing and presentation machinery. A)** Single cell RNA-sequencing of primary fetal liver erythroid progenitors reveals expression of MHC class II and associated machinery, lyso-endosomal proteases, co-stimulatory molecules, and instructive cytokines in erythroid progenitors. Data from Vanuytsel, et al. (GSE160251).

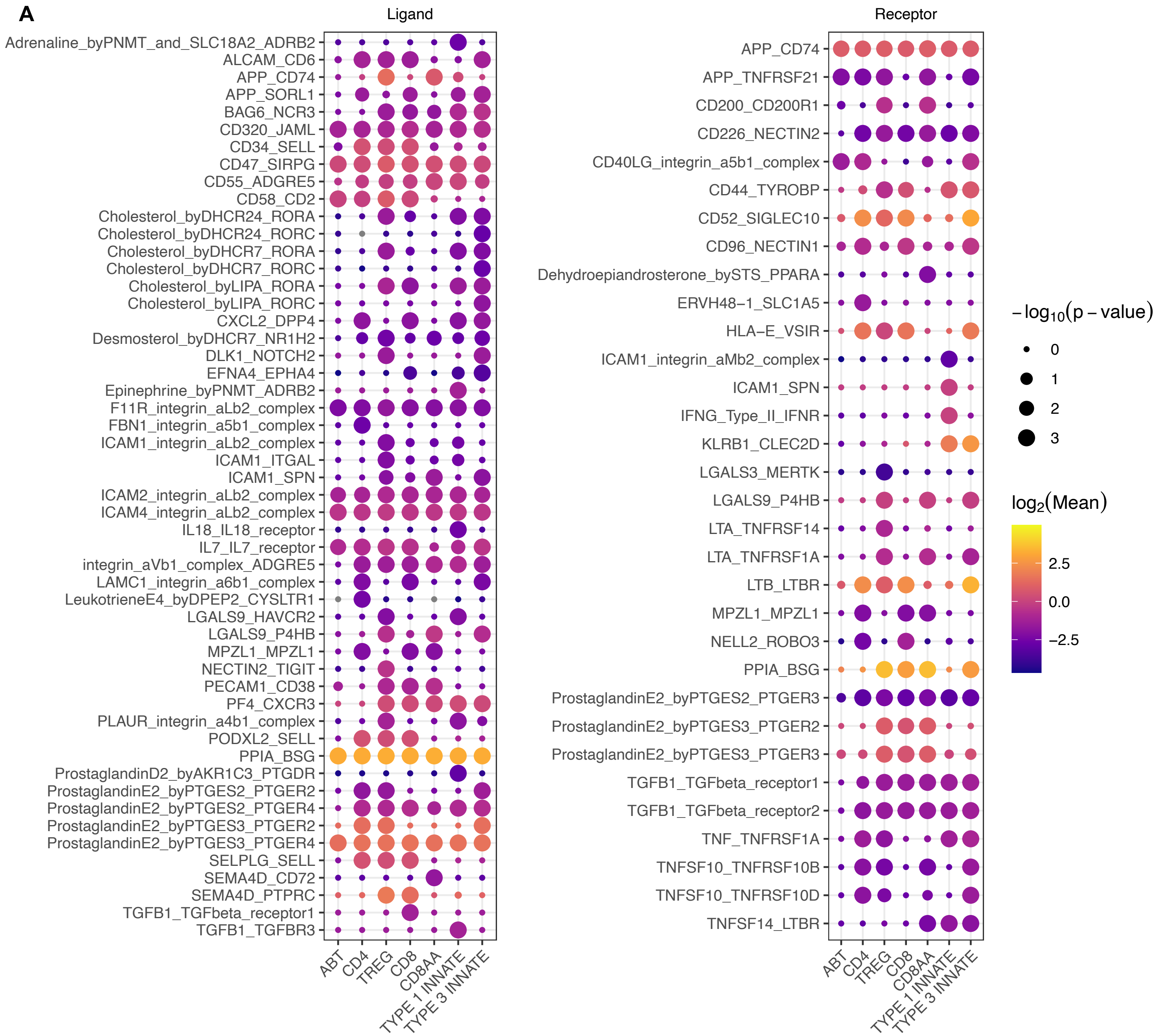


**Supplemental Figure 17. Erythroid progenitors and fetal T cells express complementary ligand-receptor pairs.** **A)** Cell-cell interactions were inferred using single cell RNA-sequencing of MHC II$^{+}$ erythroid progenitors and T cells (Data from Suo, *et al.*). Significant ligand-receptor pairs ($p<0.05$) were analyzed using CellPhoneDB. For each interaction, erythroid progenitors express either the ligand (left, first gene listed) or receptor (right, second gene listed).
